# A passage through India: the biotic ferry model supports the build-up of Indo-Australian biodiversity of an ancient soil arthropod clade

**DOI:** 10.1101/2024.05.31.596799

**Authors:** Maya Manivannan, Nehal Gurung, Gregory D. Edgecombe, Jahnavi Joshi

## Abstract

**Aim:** Scutigeromorpha is a globally distributed, ancient group of centipedes with at least 400 million years of evolutionary history. We assess the biogeographic history of the order, with a particular focus on the Peninsular Indian Plate (PIP), a Gondwanan fragment. We hypothesize that continental vicariance explains the disjunct distribution of extant scutigeromorphs, that PIP scutigeromorphs are of ancient Gondwanan origins, and that East Gondwana vicariance explains the diversification of the subfamily Thereuoneminae into its Asian and Australian clades.

**Location:** Global

**Taxa:** Centipedes, Scutigeromorpha

**Methods:** We use a novel molecular dataset sampled across the PIP and a global sequence database representing the geographic distribution of all families and subfamilies. We employ molecular phylogenetic analyses on two mitochondrial and three nuclear markers, molecular species delimitation and ancestral range estimation to reconstruct biogeographic history.

**Results:** Ancestral scutigeromorphs were likely widely distributed across Gondwana and diverged prior to any continental vicariance. Scutigeromorph biogeography is shaped by continental vicariance, long-distance dispersal and jump dispersal, indicating an ability to colonize areas far from their ancestral range. The PIP emerged as the ancestral range of Thereuoneminae, which started diversifying during the Cretaceous Period. Subsequent in-situ diversification within the PIP and dispersals into Asia, Australia, and the Pacific Islands best explained the extant distribution of Thereuoneminae, more so than East Gondwana vicariance.

**Main conclusions:** The in-situ diversification of PIP species and their ancient divergence suggest they represent Gondwanan relicts in a landmass whose biota is primarily dispersal-driven. A single dispersal event from the PIP generated most of the extant diversity in Australia, another Gondwanan fragment. Furthermore, the discovery of 11 putative species from the PIP and the Andaman Islands, five times more than was known, highlights the Wallacean and Linnean shortfalls in tropical Asia.

## Introduction

Plate tectonics has influenced species diversity and distribution patterns across different spatial scales. For instance, areas with high endemicity are associated with high tectonic activity, suggesting that the lack of gene flow between previously connected landmasses influences rates of allopatric speciation (Pellissier et al., 2017). Furthermore, tectonic movements interact with climate heterogeneity to demarcate zoogeographical boundaries (Ficetola et al., 2017). Perhaps the most well-known application of plate tectonics is in Gondwanan biogeography, with many studies investigating the relative importance of continental vicariance and dispersal in shaping the biotic composition of modern-day Gondwanan fragments (Capobianco & Friedman, 2019; Karanth, 2021; Ladiges & Cantrill, 2007; Lohman et al., 2011; Sanmartín & Ronquist, 2004; Wallis & Trewick, 2009; Yoder & Nowak, 2006). However, it is argued that signatures of vicariance may be obscured or inaccurately inferred in geologically young and vagile taxa. Therefore, groups with ancient evolutionary origins, limited dispersal ability and wide distributions are appropriate for assessing the roles of historical events in shaping extant distributions (Karanth, 2021; Kodandaramaiah, 2009).

Soil arthropods like centipedes (Class: Chilopoda) are ideal models to test Gondwanan or global vicariance biogeography hypotheses due to their high taxonomic, genetic and trait diversity and extensive evolutionary history (Bharti et al., 2023; Wong et al., 2019; Z.-Q. Zhang, 2011). The centipede order Scutigeromorpha (∼95 species) started diversifying in the Silurian Period (434 mya, 95% HPD: 412-459 mya) (Benavides et al., 2023) and has undergone little morphological diversification since then (Giribet & Edgecombe, 2013). Scutigeromorphs are readily distinguished from other centipedes by their faceted eyes, dorsally positioned spiracles, and extremely long legs that facilitate fleet locomotion; the latter is speculated to be a strategy for defense (Imms, 1910; Manton, 1965). They do not burrow deep under the soil and are often found sitting on tree bark or rock crevices or taking refuge under dead logs. It is possibly due to their agility, rarity and difficulty in morphological identification at the species level that collections and taxonomic descriptions from many parts of the world are scant, hindering the study of their evolutionary biology (Butler et al., 2010; Pérez-Gelabert & Edgecombe, 2013).

Although evolutionary relationships within the order have been comprehensively examined and dated using fossil calibrations, the underlying biogeographic history remains to be studied (Butler et al., 2010; Edgecombe & Giribet, 2006, 2009; Giribet & Edgecombe, 2013; Porta & Giribet, 2024). Scutigeromorphs have ancient divergences and largely disjunct distributions; the family Pselliodidae is Afrotropical and Neotropical in distribution, Scutigerinidae has an Afromalagasy distribution, and Scutigeridae is pantropical. The latter is further divided into subfamilies Scutigerinae, distributed in the Nearctic and Western Palearctic, and Thereuoneminae, found in the Eastern Palearctic, Oriental, and Australasian realms. This suggests the possibility of large-scale, historical vicariance events driving their global biogeography, as found in other ancient, terrestrial arthropods with vast disjunct distributions (Baker, Boyer, et al., 2020; Chamberland et al., 2022; Toussaint et al., 2017). We use quantitative historical biogeographic analyses to explicitly test continental vicariance and Gondwanan biogeographic hypotheses. Additionally, we use a novel molecular and distributional dataset of scutigeromorphs sampled across the Peninsular Indian Plate (PIP), a former Gondwanan fragment, for a complete reconstruction of biogeographical history.

The geoclimatic history of the PIP offers critical insights into the origins of tropical Asian biodiversity (Klaus et al., 2016; Kooyman et al., 2019). Originally part of the Gondwana supercontinent (∼250 mya), the PIP separated from the Australian-Antarctic region (136-126 mya) and subsequently from Madagascar (94-84 mya) and Seychelles (64 mya) after which it experienced a period of isolated rafting until its collision with the Eurasian plate approximately 59 ± 1 mya (Gibbons et al., 2013; Hu et al., 2016). Prior to the collision, the climatic regime of the PIP was frequently seasonal mega-thermal, but the uplift of the Tibetan plateau established the current Indian monsoon (Morley, 2018). Employing time-calibrated molecular phylogenies, historical biogeographic studies of soil arthropods and a few herpetofauna indicate that ancient Gondwanan taxa underwent diversification while on the drifting Indian plate and subsequently dispersed into Asia after the collision (Biju & Bossuyt, 2003; Joshi et al., 2020; Loria & Prendini, 2020; Sidharthan & Karanth, 2021). On the other hand, more recently evolved taxa dispersed into India from Central and Southeast Asia after the collision (Chen et al., 2018; Datta-Roy et al., 2012; Li et al., 2020; Sil et al., 2020). Ancient vicariance and dispersal have hence been evoked to describe the extant biogeography of the Oriental realm. However, studies using molecular phylogenies with divergence time estimates suggest that most of India’s extant tetrapod diversity was generated by post-collision dispersal events (Karanth, 2021).

We use molecular phylogenetics, molecular species delimitation analyses and ancestral range estimation analyses to test our hypotheses that 1) historical continental vicariance events drive the biogeography of Scutigeromorpha, 2) Peninsular Indian lineages are Gondwanan in origin, having diverged before the Indo-Eurasian collision and 3) the rifting of the PIP from Gondwana played a crucial role in the biogeography of Scutigeromorpha by driving the divergence of the speciose Thereuoneminae into its Asian and Australian clades. We expect that the ancestor of Thereuoneminae was distributed across East Gondwana and that continental vicariance caused the divergence of the Australian and Asian subdivisions, with the latter dispersing into Asia from the PIP post the Indo-Eurasian collision. Our results demonstrate that including PIP species is critical to inferring a complete global biogeographic history and that Gondwanan vicariance still finds its significance in a landmass whose biodiversity is primarily dispersal-driven.

## Materials and Methods

### Taxon sampling and DNA sequence generation

Scutigeromorphs were systematically sampled across the Western Ghats latitudinal gradient (n=61) and opportunistically in the Eastern Ghats (n=9) of Peninsular India between 2007-2022. We also used one specimen sampled from the Andaman Islands in 2021, totaling 71 individuals (Figure 1a, Table S1). Most of our sampling efforts were in undisturbed forested areas during the southwest (June-September) and northeast (October-November) monsoon seasons (Figure 1b and 1c). All specimens were stored in 75% ethanol at the Council of Scientific Research-Centre for Cellular and Molecular Biology (CSIR-CCMB), Hyderabad, India. We supplemented molecular data for 33 species from Giribet and Edgecombe (2013) (Table S2). This amounts to roughly 1/3rd of the global species diversity of Scutigeromorpha and samples all three families and two subfamilies. It also constitutes the largest and most comprehensive geographic sampling of the order so far, with representation across the tropics (except SE Asia) and extra- tropics (Figure 1d).

**Figure 1.**
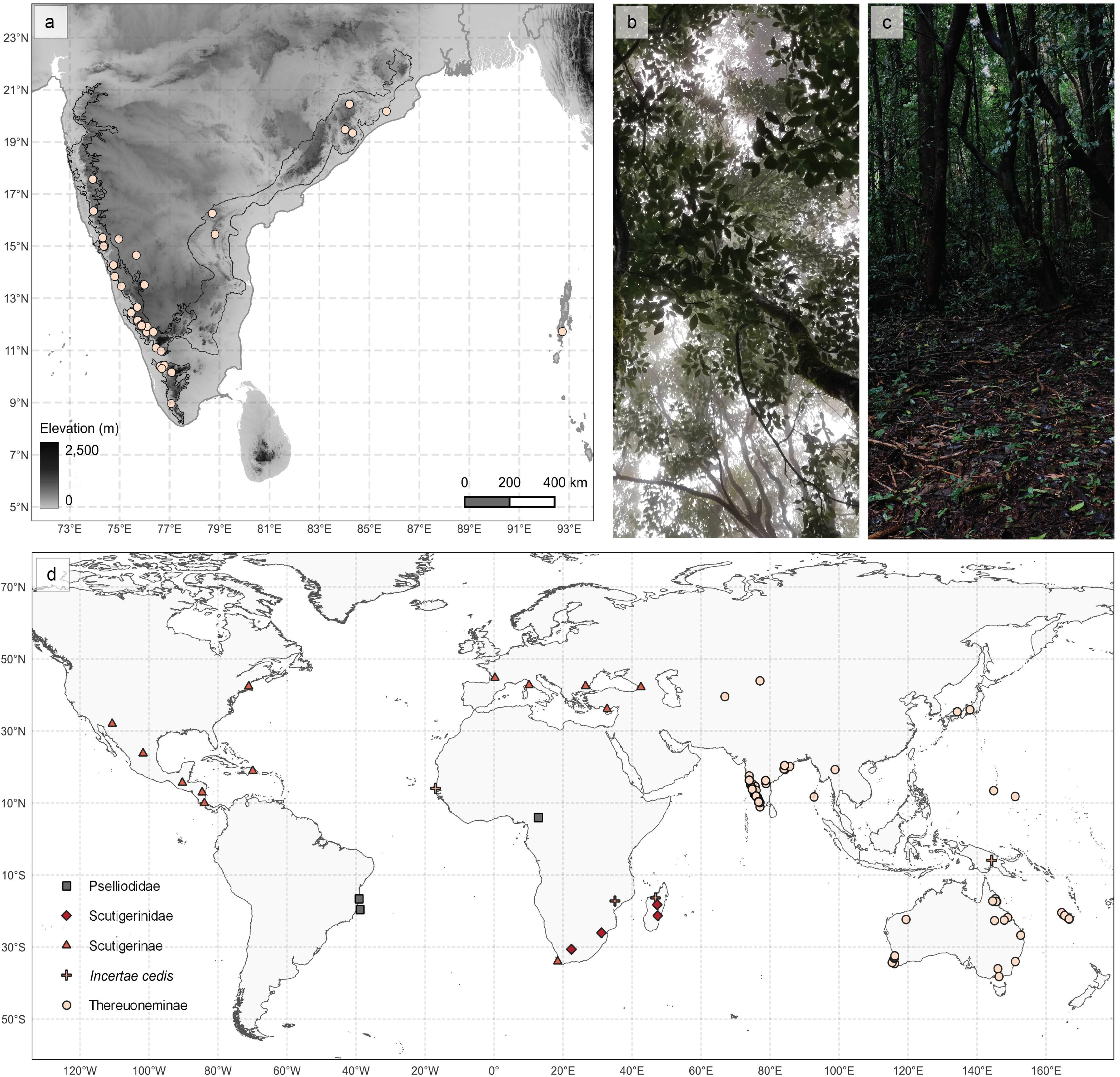
a) A map of India showing sampling locations from the Western Ghats, the Eastern Ghats and the Andaman Islands. b) and c) Examples of canopy cover and forest floor observed in collection localities in India. d) A global map representing sampling locations of all scutigeromorphs in this study sorted according to family and sub-family.

DNA extraction and PCR amplification protocols were modified from those detailed in Joshi and Karanth (2011) and Giribet and Edgecombe (2013) (Appendix 1, Table S3). PCR products were sequenced at an in-house Sanger sequencing facility at CSIR-CCMB, Hyderabad. Sequences were manually cleaned, aligned using ClustalW (Thompson et al., 1994), and concatenated in MEGA 7 (Kumar et al., 2016), totaling 2354 bp for 144 taxa (71 specimens from this study, 67 scutigeromorphs and six outgroups from Giribet and Edgecombe (2013)).

### Molecular phylogenetic analyses

We used PartitionFinder v. 1.1.1 (Lanfear et al., 2012) to identify partitioning schemes and their models of sequence evolution (MSE) under the following parameters: branchlengths = ‘linked’, models = ‘all’, model_selection = ‘AICc’, search = ‘greedy’. The five genes were identified as the best partitions. GTR+I+G and SYM+I+G were retrieved as the best-fitting MSEs for the mitochondrial and nuclear genes, respectively, and were input as such in all further analyses. Phylogenetic trees were constructed using the Maximum Likelihood (ML) and Bayesian Inference (BI) approaches. The ML tree was generated in IQTree v.1.6.12 (Nguyen et al., 2015) using partition models (Chernomor et al., 2016) with branch support values estimated using Ultrafast bootstrapping (UB2) (Hoang et al., 2018) for 1000 replicates. The BI tree was constructed with MrBayes v.3.2.7 (Ronquist et al., 2012) using parallel MCMC chains (Altekar et al., 2004) sampled every 10,000 steps for 50 million generations until the standard deviation of split frequencies dropped below 0.005. Strictly bifurcating trees were summarized using the ‘sumt contype=allcompat’ function. Log files were analyzed using Tracer v.1.7.2 (Rambaut et al., 2018) to ensure that runs converged, and all parameter ESS values exceeded 200.

### Species delimitation analyses

To arrive at a putative species list of scutigeromorphs newly sampled from India, we employed two single-locus heuristic methods of species delimitation- Assemble Species by Automatic Partitioning (ASAP, Puillandre et al., 2021) and Poisson Tree Processes (PTP, J. Zhang et al., 2013). We delimited species using the COI and 16S datasets because of sufficient sequence variability (49% and 69%, respectively) and low effective population size (mitochondrial markers). We evaluated the different delimitation methods based on result convergence, well-supported monophyly of delimited species in ML and BI phylogenies and success in delimiting species established in literature according to morphological and molecular data. For ASAP, FASTA files were analyzed on the web server (https://bioinfo.mnhn.fr/abi/public/asap/<u>#</u>) (Puillandre et al., 2021) using default settings (JC69 substitution model, 0.01 split group probability). Input phylograms for PTP were created with MrBayes v.3.2.7 (Ronquist et al., 2012) using parallel MCMC chains (Altekar et al., 2004) for 50 million generations sampled every 10,000 steps and verified for convergence on Tracer v.1.7.2 (Rambaut et al., 2018). The analysis was run on the mPTP linux tool (Kapli et al., 2017) (‘--single’) after cropping outgroups (--outgroup_crop) and calculating the minimum branch length (--minbr_auto). MCMC was run with four chains for 10 million generations, a sampling frequency of 10,000 and a burn-in of 10% along with commands ‘--mcmc_log’ and ‘--mcmc_startnull’.

### Divergence time estimation

A time-calibrated phylogeny was generated in StarBEAST2 (Ogilvie et al., 2017), which offers a computationally efficient, multi-species coalescent approach to inferring species trees from concatenated sequence alignments. Trees, clocks and site models were linked across the mitochondrial genes but independently analyzed for the nuclear genes. A relaxed lognormal clock model (Drummond et al., 2006) was used with a constant population function and a Yule speciation prior. MRCA (Most Recent Common Ancestor) priors were included in the analysis to constrain taxon relationships according to those obtained from the ML and BI phylogenies. We used five fossil calibrations from Wolfe et al. (2016) to estimate divergence times (Table S4). Uniform distribution priors were selected for all calibration points. The analysis was run for 1 billion generations and sampled every 1000 trees. Convergence was verified in Tracer v.1.7.2 (Rambaut et al., 2018). Trees were summarized as Maximum Clade Credibility (MCC) trees with median node heights after discarding a burn-in of 50% on TreeAnnotator (Drummond & Rambaut, 2007).

### Ancestral range estimation

Distributions of scutigeromorph species represented in this study were identified using ChiloBase 2.0 and prior literature for molecular voucher specimens (Bonato et al., 2016; Edgecombe & Giribet, 2006, 2009; Giribet & Edgecombe, 2013). Singleton species from Giribet and Edgecombe (2013) that are yet to be described were assigned their sampling locations. In total, 11 areas were defined: Africa (African continent with the Arabian Peninsula), Australasia (the Sahul region consisting of Australia and Papua New Guinea), Madagascar, Pacific Islands (Guam and the Federated States of Micronesia), Nearctic, Neotropics, New Caledonia, Palearctic (Palearctic biogeographic zone excluding North Africa and the Arabian peninsula), Peninsular Indian Plate (present-day Indian peninsula along with Sri Lanka, parts of Pakistan and Bangladesh), SE Asia (continental SE Asia along with western Malesia that was part of Gondwana; referred to as the Cimmerian terrane) and the Andaman and Nicobar islands. Other prominent areas, such as the rest of insular SE Asia and New Zealand, were not coded as they lacked sufficient representation.

The global biogeographic history of Scutigeromorpha was assessed with BioGeoBEARS implemented in R v.4.3.1 (Matzke, 2018; R Core Team, 2023). We employed the DEC (Dispersal-Extinction-Cladogenesis) model and the maximum likelihood versions of DIVA (Dispersal-Vicariance Analysis) and BAYAREA (DIVALIKE and BAYAREALIKE respectively), along with their ‘+j’ (founder-event speciation) versions (Landis et al., 2013; Matzke, 2013, 2014; Ree et al., 2005; Ree & Smith, 2008; Ronquist, 1997). Including founder-event speciation, which has been argued to play an important role in the biogeographic history of several taxa, improves fit when taxa mostly occupy single-area ranges (Matzke, 2014, 2022).

The biogeographic models were implemented in three time-stratified schemes: 1) constraining areas allowed and between-area dispersal probabilities, 2) no constraint on areas allowed but constraining between-area dispersal probabilities and 3) constraining areas allowed but no constraints on between-area dispersal probabilities. Model fits were compared using the Akaike Information Criterion corrected for small sample sizes (AICc). Other models (non-stratified, no constraint on areas allowed or between-area dispersal probabilities) were not considered as the lack of tectonic or ecological information renders them too simplistic.

Five-time points representing main tectonic activity were chosen for the time-stratified analysis: 150 mya (rifting between East and West Gondwana), 120 mya (rifting between PIP and the rest of East Gondwana), 100 mya (rifting between the Neotropics and Africa), 40 mya (collision between PIP and the Palearctic, the emergence of New Caledonia and Guam) and 15 mya (emergence of the Andaman and Nicobar Islands) (De Bruyn et al., 2014; Grandcolas et al., 2008; Matthews et al., 2016; Müller et al., 2016; Neall & Trewick, 2008). Dispersal matrices were created by referencing the above literature (Table S5). The maximum number of areas in a range was constrained to three. Analyses constrained with an ‘Areas allowed’ text ensured that some of the islands (Pacific Islands, New Caledonia, Andaman and Nicobar) were excluded from the analysis at time points before their origin. We supplemented the ancestral range estimation analyses with Biogeographic Stochastic Mapping (BSM) implemented in BioGeoBEARS for 100 iterations to estimate the probability and type of biogeographic events at nodes for which range resolution was poor (Dupin et al., 2017; Matzke, 2018; R Core Team, 2023).

## Results

### Molecular phylogeny and species delimitation

The tree topologies from the ML and BI approaches were highly congruent, and most relationships were well-supported (posterior probability > 0.85 / bootstrap support > 95) (Supplementary figure). Our phylogeny closely resembled that obtained by Giribet and Edgecombe (2013), with two major exceptions: Scutigerinae (a subfamily of Scutigeridae) was paraphyletic rather than monophyletic, and the *Incertae cedis* group comprised of *Ballonema* and *Lassophora* was monophyletic (1/98).

Scutigeromorphs from the Indian subcontinent are part of the subfamily Thereuoneminae within Scutigeridae. Clade 1, consisting of members of the genus *Thereuopoda,* was strongly supported (0.93/100) and was a sister group to *Thereuopoda longicornis* from Southeast Asia (1/100). An individual (CCMB10) from the Andaman Islands was sister to *Thereuopodina* sp. from Australia with low support (0.72/84). This clade was sister to an individual from the Eastern Ghats (CCMB4071) with high support (0.91/94). Clade 2, monophyly of which was moderately supported (0.71/96). was sister to (CCMB4071, (*Thereuopodina* sp., CCMB10)) with low support (0.53/79). Clade 4 (0.66/82) was sister to the Australia-New Caledonia genera *Allothereua-Parascutigera-Pilbarascutigera* with low support (0.69/85). However, the sister relationship between the two and Thereuoneminae spp. from the Pacific Islands was well supported (1/100). Clade 3 (1/100) was sister to (Thereuoneminae sp., (*Allothereua- Parascutigera-Pilbarascutigera*, Clade 4)) with low support (0.51/80).

We reconciled the results from two single-locus species delimitation methods, ASAP and PTP, to get a conservative estimate of the number of Indian species based on convergence between species delimitation analyses, sequence data availability and monophyly for Indian clades (Figure 2, Appendix 2, Table S6). The same procedure confirmed species identities for scutigeromorphs from the global dataset. Species delimitation analyses, monophyly, and geographic distribution indicated at least 11 species in our dataset across four clades in the PIP.

**Figure 2.**
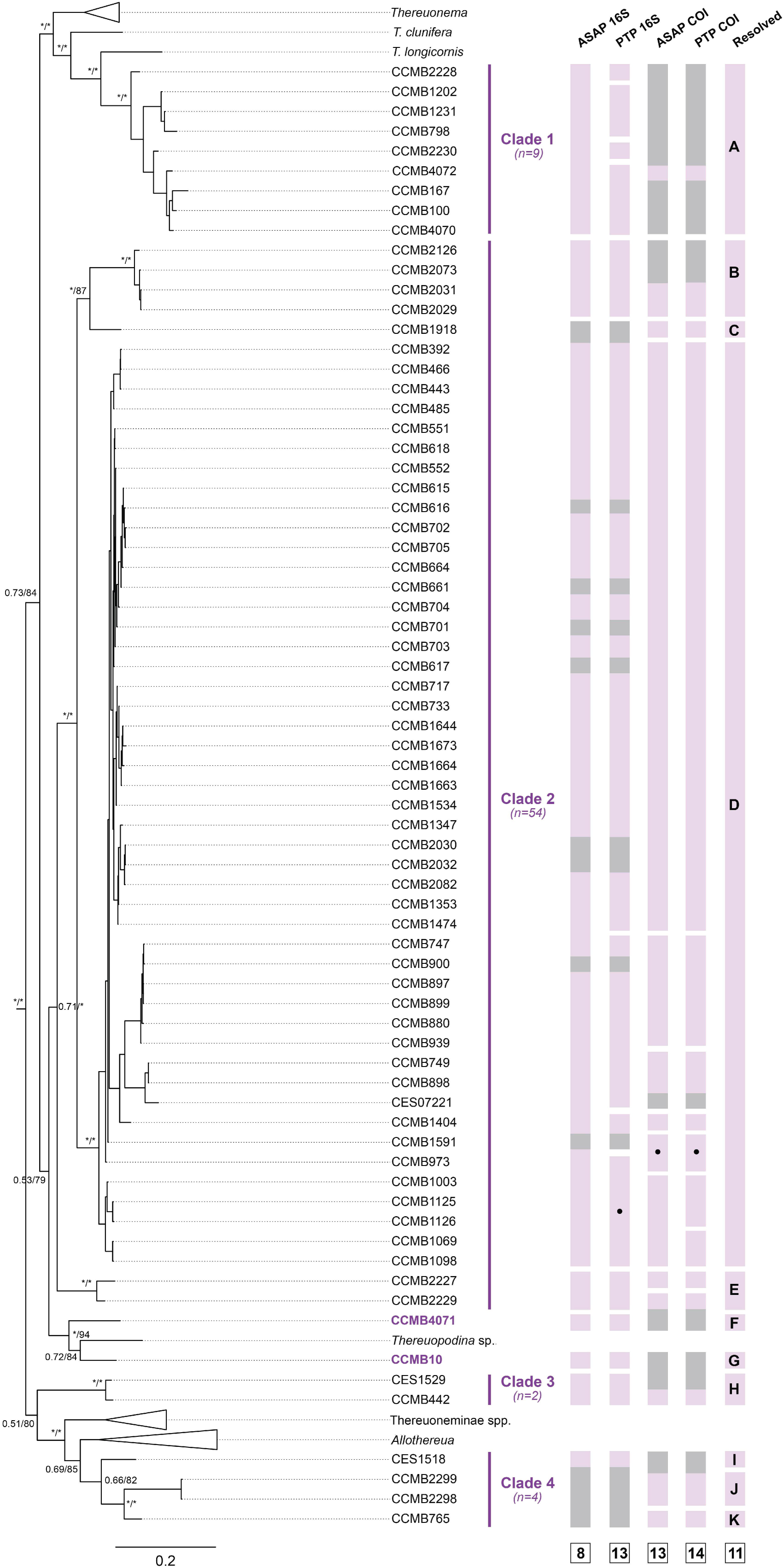
Species delimitation results for Scutigeromorphs sampled in India. Left-expanded molecular phylogeny of Scutigeromorpha from concatenated COI, 16S, 28S, H3 and 18S genetic data. Bootstrap supports are indicated at nodes as BI/ML. Asterisks indicate support values > 0.85 for Bayesian analyses and > 95 for UB2 in Likelihood analyses. Right-Species identities from ASAP and PTP on COI and 16S data. Each bar in a column indicates a unique species. Bars with a circle are not unique species but are part of a larger delimited species group. Gray regions represent missing genetic data. The last column refers to resolved species identities used for downstream analyses, allocated with species names A-K.

### Divergence times and biogeography analyses

From the ancestral range estimation, two models, DEC+J constrained with a variable dispersal matrix and areas allowed (lnL -111.7) and DEC+J constrained with a variable dispersal matrix but with no constraint on areas allowed (lnL -111.4), had the lowest AICc scores (Table 1). The ancestral area reconstructions of these two models were identical; hence, we present the results of model 1 for subsequent discussion along with BSM probabilities. Two vicariance, six dispersal, and four jump dispersal events best explained the global distribution of Scutigeromorpha (Figure 3). The first split within Scutigeromorpha was into Pselliodidae, and the lineage comprised of Scutigeridae and Scutigerinidae at the boundary of the Devonian and Carboniferous (354 mya; 95% HPD: 237-466 mya). The ancestral ranges of some of the deeper nodes at the family level were uncertain, with low probability values for multiple possible ranges. BSM allowed us to incorporate uncertainty in their range estimation, indicating that ancestral scutigeromorphs were most likely distributed only in Gondwanan areas, as shown in Figure 3. Furthermore, from the origin of Scutigeromorpha until the divergence of *Tachythereua*+*Scutigera* from the rest of Scutigeridae, the ranges of ancestral scutigeromorphs most likely occupied more than 1 Gondwanan area. BSM results also supported the role of subset sympatry in the diversification of several lineages, where one daughter clade acquired the ancestral range, and the other acquired only a part of the range wherein the two clades were sympatric (see Table 2 for a summary of cladogenetic events at poorly resolved nodes).

**Figure 3.**
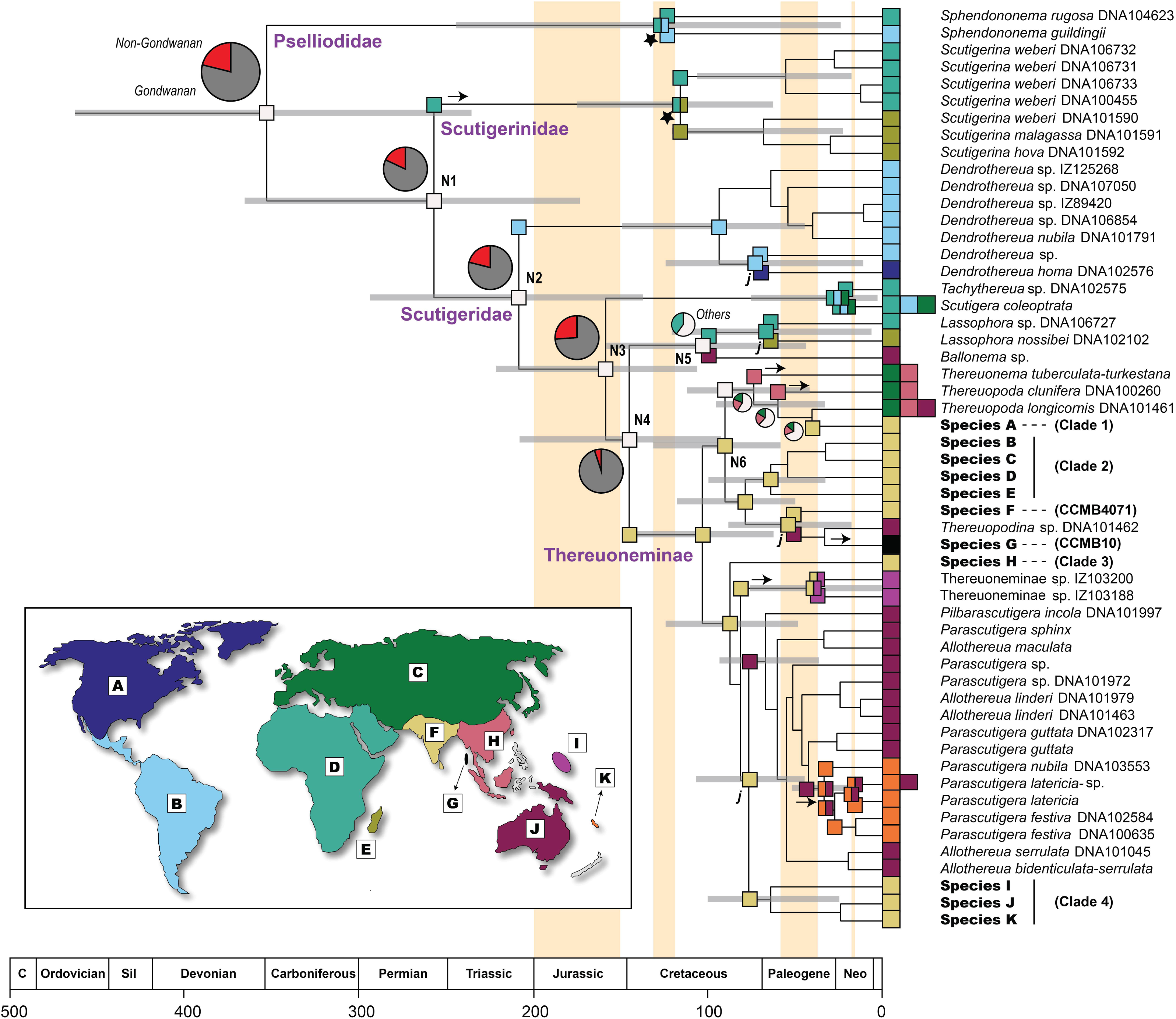
Ancestral range estimation for dated phylogeny of Scutigeromorpha. Gray bars at nodes correspond to 95% HPD credible intervals for node ages. Coloured boxes at nodes and tips represent areas as indicated in the inset map (A: Nearctic, B: Neotropics, C: Palearctic, D: Africa, E: Madagascar, F: Peninsular Indian Plate, G: Andaman Islands, H: Southeast Asia, I: Pacific Islands, J: Australia, K: New Caledonia). Only ancestral ranges with ≥ 50% probability according to best-fitting DEC+J models are recorded. Nodes with poor resolution (< 50%) are represented in white with pie charts indicating proportions of ancestral ranges. Nodes labeled N1-N6 are discussed further in the text. Anagenetic dispersal events are represented as arrows. Cladogenetic events are represented as stars for vicariance and ‘j’ for founder-event speciation/jump dispersal. Pale yellow bars behind phylogeny indicate the following geological events: split between East and West Gondwana (200-150 mya), split between PIP and East Gondwana (130-120 mya), collision between Peninsular Indian and Eurasian plates (60-40 mya) and origin of Andaman Islands (15 mya).

**Table 1.** Scores for BioGeoBEARS models. Log likelihoods, number of parameters, parameter values for dispersal, extinction and jump dispersal, Akaike Information Criterion corrected for small sample sizes (AICc), delta AICc and AICc weights are indicated. The rows highlighted in bold correspond to the two models with the lowest AICc scores.

| Model | LnL | N | d | e | j | AICc | Delta AICc | AICc weight |
| --- | --- | --- | --- | --- | --- | --- | --- | --- |
| **1: Constraining areas allowed and between-area dispersal probabilities** |  |  |  |  |  |  |  |  |
| DEC | -115.7 | 2 | 0.0014 | 0.0012 | 0 | 235.7 | 6.3 | 0.022 |
| **DEC + J** | **-111.7** | **3** | **0.0009** | **0.0008** | **0.034** | **229.9** | **0.5** | **0.408** |
| DIVALIKE | -122.3 | 2 | 0.0018 | 0.0014 | 0 | 248.8 | 19.4 | 3.20E-05 |
| DIVALIKE + J | -118.3 | 3 | 0.0011 | 0.0009 | 0.037 | 243 | 13.6 | 0.0006 |
| BAYAREALIKE | -129.5 | 2 | 0.0016 | 0.0075 | 0 | 263.3 | 33.9 | 2.28E-08 |
| BAYAREALIKE+J | -118 | 3 | 0.0005 | 0.0014 | 0.064 | 242.5 | 13.1 | 0.0007 |
| **2: No constraint on areas allowed but constraining between-area dispersal probabilities** |  |  |  |  |  |  |  |  |
| DEC | -115.2 | 2 | 0.0014 | 0.001 | 0 | 234.5 | 5.1 | 0.0409 |
| **DEC + J** | **-111.4** | **3** | **0.0009** | **0.0007** | **0.031** | **229.4** | **0** | **0.523** |
| DIVALIKE | -122.1 | 2 | 0.0018 | 0.0014 | 0 | 248.5 | 19.1 | 3.73E-05 |
| DIVALIKE + J | -118.1 | 3 | 0.0011 | 0.0009 | 0.041 | 242.7 | 13.3 | 0.0007 |
| BAYAREALIKE | -129.2 | 2 | 0.0016 | 0.0075 | 0 | 262.6 | 33.2 | 3.23E-08 |
| BAYAREALIKE+J | -117.7 | 3 | 0.0005 | 0.0015 | 0.064 | 241.9 | 12.5 | 0.00101 |
| **3: Constraining areas allowed but no constraint on between-area dispersal probabilities** |  |  |  |  |  |  |  |  |
| DEC | -121.1 | 2 | 0.0006 | 0.0012 | 0 | 246.5 | 17.1 | 0.0001 |
| DEC + J | -116.8 | 3 | 0.0004 | 0.0007 | 0.014 | 240.1 | 10.7 | 0.00249 |
| DIVALIKE | -128.1 | 2 | 0.0007 | 0.0013 | 0 | 260.3 | 30.9 | 1.02E-07 |
| DIVALIKE + J | -124.6 | 3 | 0.0005 | 0.001 | 0.013 | 255.6 | 26.2 | 1.07E-06 |
| BAYAREALIKE | -134.3 | 2 | 0.0006 | 0.0067 | 0 | 272.9 | 43.5 | 1.87E-10 |
| BAYAREALIKE+J | -121.8 | 3 | 0.0002 | 0.0023 | 0.018 | 250.1 | 20.7 | 1.67E-05 |

**Table 2.**
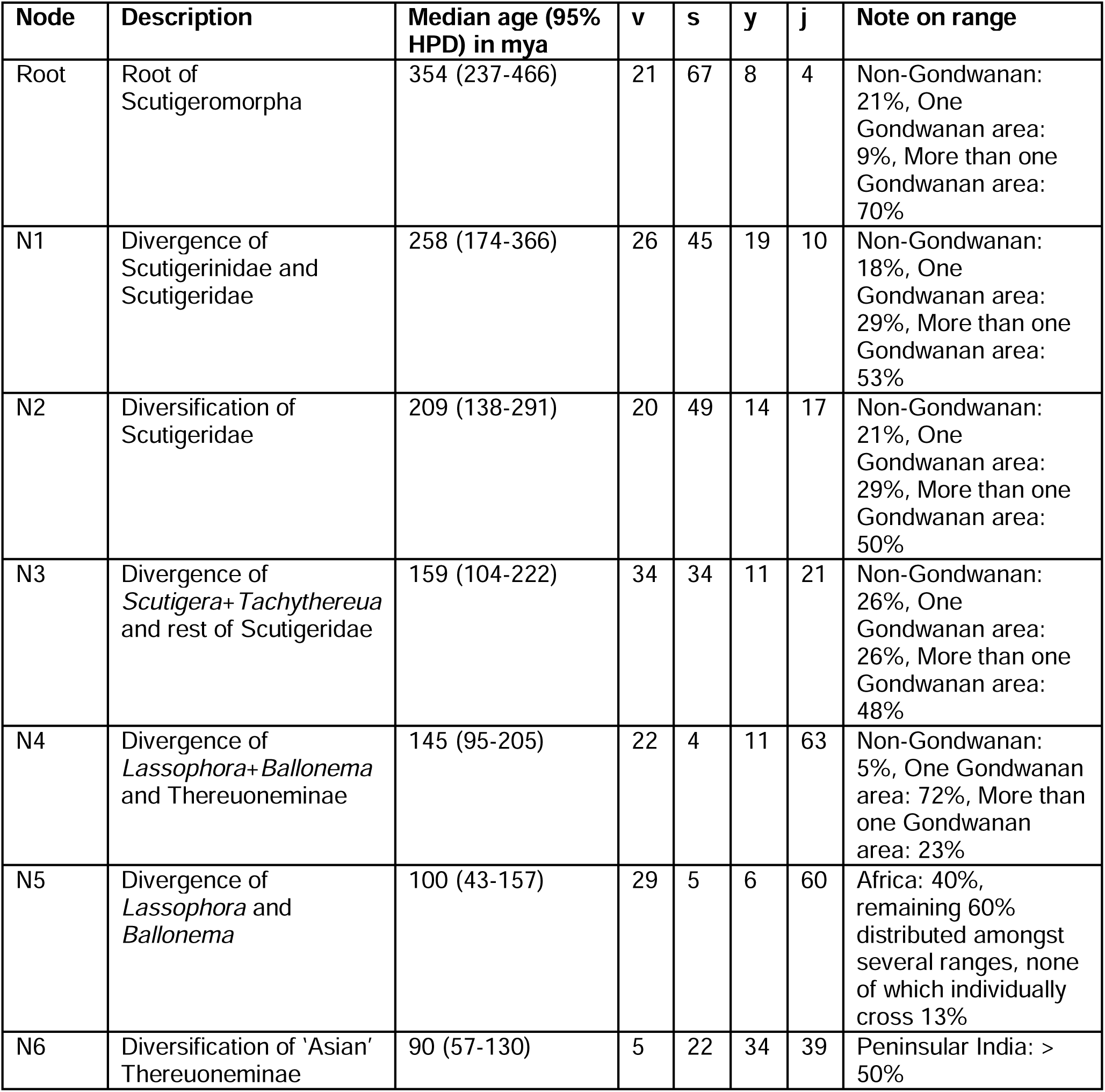
Percentages of cladogenetic events (vicariance, subset sympatry, sympatry and jump dispersal) at nodes with low ancestral range resolution estimated by Biogeographic Stochastic Mapping. Nodes indicated as per Figure 3.

The family Pselliodidae continued to diversify, and African *Sphendononema rugosa* and Neotropical *S. guildingii* split around 124 mya (95% HPD: 27-245 mya) due to a vicariance event. The other two families, Scutigerinidae and Scutigeridae, diverged from each other with a midpoint dated to 258 mya (95% HPD: 174-366 mya). The ancestor of Scutigerinidae dispersed from Africa to Madagascar, following which an Africa-Madagascar vicariance caused the divergence between the African and Malagasy *Scutigerina* around 116 mya (95% HPD: 65-178 mya). The earliest split in the Scutigeridae family was around 209 mya (95% HPD:138-291 mya). The predominantly Neotropical *Dendrothereua* started diversifying in the Late Cretaceous around 93 mya (95% HPD: 42-151 mya) and had one instance of jump dispersal into the Nearctic near the boundary of the Cretaceous and Paleogene around 69 mya (95% HPD:12- 127 mya). A subset sympatry event occurred at the ancestral node of *Tachythereua* sp. and widely distributed *Scutigera coleoptrata* between Africa+Neotropics+Palearctic and Africa very recently at the Paleogene-Neogene boundary around 21 mya (95% HPD: 2-75 mya). Further within Scutigeridae, there was a possible jump dispersal event around 145 mya (95% HPD: 95- 205 mya) from a single Gondwanan area, leading to the clade of *Lassophora* and *Ballonema* in Africa, Madagascar and Australasia and subfamily Thereuoneminae found predominantly in the Asian-Australian Region. Within the *Incertae cedis* clade of *Lassophora* and *Ballonema*, the divergence of *Ballonema* from *Lassophora* was likely due to a jump dispersal event from Africa to Australia approximately 100 mya (95% HPD: 43-157 mya). A subsequent jump dispersal event was recovered at the ancestral node for *Lassophora*, from Africa to Madagascar, leading to *L. nossibei* at the beginning of the Paleogene at 64 mya (95% HPD: 6-120 mya).

Thereuoneminae started diversifying around 103 mya (95% HPD: 64-144 mya), and the PIP was recovered as the ancestral area. One clade of Thereuoneminae was composed mostly of Asian species, whereas the other was primarily Australian, with a few PIP species. Morphology-based taxonomic data suggest that the latter clade includes *Allothereua wilsonae* from Nepal (Dobroruka, 1978) and *Allothereua kirigisorum* from Kazakhstan (Lignau, 1929). The PIP was recovered as the ancestral range for both daughter clades.

The Asian Thereuoneminae clade started diversifying around 90 mya (95% HPD: 57-130 mya). Although the ancestral range of one of its daughter clades (containing Indian Species B-G and *Thereuopodina* sp. from Australia) was resolved as the PIP, the ancestral range of the other daughter clade (*Thereuonema-Thereuopoda* and PIP Species *Thereuopoda* sp.) was unclear, preventing us from commenting on the biogeographical event behind this divergence. Subsequent dispersal events within the latter clade into the Palearctic from Southeast Asia (leading to *Thereuonema* and *Thereuopoda clunifera* between the last 33-111 my) were well resolved. The clade containing Species B-G and *Thereuopodina* sp. diversified through the Cretaceous (78 mya; 95% HPD: 48-116 mya). One jump dispersal event from the PIP to Australia was recovered around 51 mya (95% HPD: 20-87 mya). Following this, a dispersal event from Australia to the Andaman Islands likely followed by extinction in Australia was resolved where Species G diverged 33 mya (95% HPD: 0-66 mya) from Australian *Thereuopodina* sp.

The Australian Thereuoneminae clade began diversifying around 87 mya (95% HPD: 51-123 mya) when Species H diverged from the rest of the clade. Further, within the clade, a dispersal event from the PIP to the Pacific Islands within the last 114 mya, followed by a subset sympatry event between Pacific Islands-PIP and PIP approximately 37 mya (95% HPD: 0-73 mya), led to the Thereuoneminae spp. clade. A jump dispersal event from the PIP to Australia at approximately 76 mya (95% HPD: 44-107 mya) resulted in the divergence of the Australian-New Caledonian *Allothereua-Parascutigera-Pilbarascutigera* from PIP Species I-K. The latter clade began diversifying around 67 mya (95% HPD: 38-93 mya). A dispersal event from Australia into New Caledonia, followed by three instances of subset-sympatry between Australia-NC and New Caledonia, resulted in the divergence of New Caledonian lineages from Australian lineages and their subsequent diversification during the Oligocene-Miocene.

## Discussion

The biogeography of ancient scutigeromorph centipedes was shaped by historical continental shifts, climate-driven vicariance events, and jump and long-distance dispersals. Notably, the ancestor of the Indo-Australian Thereuoneminae, which diversified during the Cretaceous, was distributed on the Peninsular Indian Plate, a Gondwanan fragment, indicating its Gondwanan origin. In-situ diversification on the PIP, along with dispersals into Asia, Australia, and the Pacific Islands, played significant roles in shaping the current distribution of Thereuoneminae, rather than vicariance events in East Gondwana. In particular, a single dispersal event contributed to most of the extant diversity in Australia. We also propose a robust species hypothesis using molecular species delimitation methods for global scutigeromorphs, identifying 11 putative species from the PIP and the Andaman Islands. Previous research had suggested the distribution of only two species in the PIP, highlighting gaps in taxonomic and distributional data for soil arthropods in the region, Linnaean and Wallacean shortfalls, respectively (Hortal et al., 2015).

### Phylogenetics and systematic implications

Relationships within Scutigeromorpha are similar to previous studies (Butler et al., 2010; Edgecombe & Giribet, 2006, 2009; Giribet & Edgecombe, 2013). Scutigeromorphs from the Peninsular Indian Plate are part of the Asian and Australian Thereuoneminae clades, as expected from distribution patterns (Edgecombe & Giribet, 2006, 2009; Giribet & Edgecombe, 2013). Two species of Scutigeromorpha have been previously reported from the PIP: *Thereuopoda longicornis* (Krishnan & Prasad, 2022; Würmli, 1979), which has a wide distribution across the Australasian and Oriental regions, and *Thereuopodina adjutrix* (Verhoeff, 1936), whose type locality is in the PIP. Species delimitation analyses with conservative estimates suggest that at least 11 species exist in Peninsular India and the Andaman Islands. One species (*Thereuopoda sp.,* Species A) is well-supported and is a sister to *T. longicornis*. Some material from the recent morphological examination of *T. longicornis* from the Western Ghats, India (Krishnan & Prasad, 2022) is likely *Thereuopoda sp.* (Species A) and not *T. longicornis* from Asia. Species B-G may belong to the genus *Thereuopodina,* which currently contains only three known species: *T. queenslandica* (type specimen from Australia), *T. tenuicornis* (type specimen from Sri Lanka), and *T. adjutrix* (type specimen from India). It is an exciting possibility that one of our sampled species is a rediscovery of *T. adjutrix*, 88 years after its description. Although the holotype of *T. adjutrix* has been illustrated in modern times (Unsöld & Melzer, 2003), its identification is complicated by its cited type locality (Madras), likely the port from which it was exported rather than a collecting locality. Morphological data will be needed to determine the generic identities of the four remaining taxa, Species H-K, which are sister to predominantly Australian and New Caledonian endemics and may represent novel genera altogether.

While we established a robust species hypothesis for scutigeromorphs for our global biogeographic analysis, it’s imperative that we undertake morphology-based systematic research on PIP species in the future. However, it is worth noting that identifying scutigeromorphs at the species level is notoriously difficult (see Edgecombe, 2007; Edgecombe & Barrow, 2007). Historically, the clade’s taxonomy went from an era of naming many species based on minor variations in a suite of meristic characters (e.g., numbers of articles in the flagella of the antennae and tarsomeres in the legs) but with access to only small sample sizes, to an era of rampant synonymy. Molecular species delimitation methods provide an independent means of asking interesting, large-scale questions in biogeography. It also facilitates systematics research in understudied taxa and unexplored areas like the PIP (Joshi & Agarwal, 2021). Our study indicates that the scutigeromorph diversity of the PIP is five times more than previously established.

### Continental and climatic vicariance and long-distance dispersals shape biogeography

The earliest divergences in Scutigeromorpha (during the Palaeozoic 237-466 mya) precede any Gondwanan rifting by at least 30 my (Matthews et al., 2016; Müller et al., 2016), although some overlap with initial Pangaean rifting (∼240 mya, see Matthews et al., 2016). Furthermore, many deep nodes occupy more than one Gondwanan area, with subsequent diversification occurring via subset sympatry. Our biogeography reconstruction suggests that ancestral scutigeromorphs were widely distributed in Gondwana and that their initial diversification occurred before any continental vicariance events. Other ancient taxa, such as harvestmen and velvet worms, also show similar patterns (Baker, Boyer, et al., 2020; Baker, Sheridan, et al., 2020; Murienne et al., 2014). Vicariance due to climatic disruption or dispersal mediated by climatic bridges is a possible alternative, as Pangaea and Gondwana exhibited diverse climates through geological time (Li et al., 2020; Rossini et al., 2022). For instance, Heavens et al. (2015) find evidence for high variability in precipitation within equatorial Pangaea during both glacial and interglacial cycles within the late Palaeozoic ice age. However, the need for sufficient geoclimatic resolution at such ancient timescales may make it challenging to test such climatic hypotheses explicitly. Notably, soil arthropods may have survived extreme climatic events such as the K-Pg extinction due to low body size and sheltering ability, supporting their ancient Gondwanan origins (Robertson et al., 2004). The few Indian taxa recovered as Gondwanan entities are indeed soil arthropods and burrowing herpetofaunal lineages (Biju & Bossuyt, 2003; Joshi et al., 2020; Loria & Prendini, 2020; Sidharthan & Karanth, 2021).

We find evidence for two different continental vicariance scenarios; one is the divergence of Pselliodidae into its African and Neotropical lineages (27-245 mya), which overlaps with the rifting between Africa and the Neotropics approximately 140-100 mya (Matthews et al., 2016; Müller et al., 2016), supporting West-Gondwana vicariance. Another is the diversification of Scutigerinidae into its African and Malagasy clades (65-178 mya), which overlaps with the Africa-Madagascar rift between 180 and 160 mya (Müller et al., 2016). Additionally, some long-distance jump dispersal and anagenetic dispersal (range expansion) events are recovered in the biogeographic analyses, indicating a capacity for scutigeromorphs to survive and successfully colonize areas far from their original ranges. For example, the divergence between the African and Malagasy lineages of *Lassophora* is supported by a jump dispersal from Africa to Madagascar after the rifting of the two landmasses (160-180 mya, see Müller et al., 2016), possibly facilitated by ephemeral land bridges between the Late Cretaceous and the present (Masters et al., 2021), contradicting the earlier proposed vicariance scenario (Giribet & Edgecombe, 2013). The New Caledonian radiation within sub-family Thereuoneminae (17-51 mya) is due to dispersal into New Caledonia from Australia, followed by three subset-sympatry events between NC and Australia-NC. The complex geology of the Zealandia region could have facilitated these divergences, where land continuity between Australia, New Zealand, and New Caledonia fluctuated from the Early Paleocene to the Late Eocene (approximately 40-65 mya) (Ladiges & Cantrill, 2007).

A Cretaceous-Paleogene dispersal event from PIP to PIP+Pacific Islands followed by a subset-sympatry event leads to the diversification of Thereuoneminae spp. from Guam and Micronesia. Not only is the divergence of Thereuoneminae spp. from its sister group sufficiently ancient (47- 114 mya) to negate human-mediated dispersal, but specimens from the two islands are different species according to molecular species delimitation. A similar scenario was observed in the arachnid order Schizomida, where specimens from different islands in the Pacific Islands were different species despite having originated from a single recent dispersal event (within the last 70 my), suggesting high dispersal ability coupled with insular range-restriction (Clouse et al., 2017). This indicates the possibility of common dispersal routes for taxa from Asia to the remote islands of the Pacific, perhaps via favorable ocean currents such as the North Equatorial/North Pacific current (see Demeulenaere & Ickert-Bond, 2022 for a comprehensive review of the origins of Micronesian biota).

### Ancient Gondwanan scutigeromorphs of the Peninsular Indian Plate

The Peninsular Indian Plate alone was ancestral for Thereuoneminae during the Cretaceous Period (64-144 mya) instead of PIP+Australia, as the East Gondwana vicariance hypothesis predicted. The ancient divergence date, Gondwanan ancestral distribution, and primarily Gondwanan distribution of extant scutigeromorphs indicate that Thereuoneminae is a Gondwanan group. PIP is also retrieved as the ancestral range for the Asian and Australian clades. The ancestor of the Asian clade diversified into the predominantly Southeast Asian- Palearctic genera *Thereuonema* and *Thereuopoda* and the majority of the PIP species sometime between 57 and 130 mya. The credible interval for this divergence overlaps with the Indo-Eurasian collision timing, suggesting Out-of-India dispersal into Southeast Asia. It could also support the Burma-Terrane (BT) hypothesis, where the PIP was contiguous with the Burma Terrane until about 75 mya, with the BT colliding with SE Asia at approximately 38 mya (Bolotov et al., 2022). It is possible that the divergence between the *Thereuonema-Thereuopoda* group and the PIP sister group occurred due to the rifting of the PIP-BT landmasses, with the former carried to Southeast Asia via the Burma Terrane. We hope that further sampling in Southeast Asia will enable testing this scenario.

At the ancestral node of (Species F, (*Thereuopodina* sp., Species G) within the Asian clade, as well as at the ancestral node of PIP Clade 4 (consisting of Species I, J and K) and the Australian-New Caledonian radiation of *Allothereua*, *Parascutigera* and *Pilbarascutigera* within the Australian clade, a jump dispersal from the PIP to Australia is recovered. These events occur roughly within the same period (20-87 mya and 44-107 mya, respectively) and overlap with the Indo-Eurasian collision, providing evidence for Out-of-India dispersal into Australia, although the route- via transoceanic or stepping-stone dispersal through Southeast Asia- remains unclear, and requires additional genetic representation from SE Asia to verify. Many PIP species resulted from in-situ diversification on the Indian plate, possibly representing Gondwanan relics. The lone species from the Andaman Islands (Species G) is sister to *Thereuopodina* sp. from Australia and likely dispersed from Australia within the last 66 million years, followed by extinction in Australia. This needs further verification with more sampling in the Andaman and Nicobar Islands and Southeast Asia.

### Biogeographic relationships between Australia and Peninsular India

Species from the Peninsular Indian Plate were part of two distinct clades with Asian and Australian affinities within Thereuoneminae. Sister or nested relationships between Asian and PIP taxa are well-documented and explained by the ‘Out-of-India’ and ‘Into-India’ hypotheses (Datta-Roy & Praveen Karanth, 2009). However, instances of PIP-Australia relationships are much rarer. Here, we synthesize biogeographic studies to compare our results with a few empirical examples showcasing such a pattern (Figure 4).

**Figure 4.**
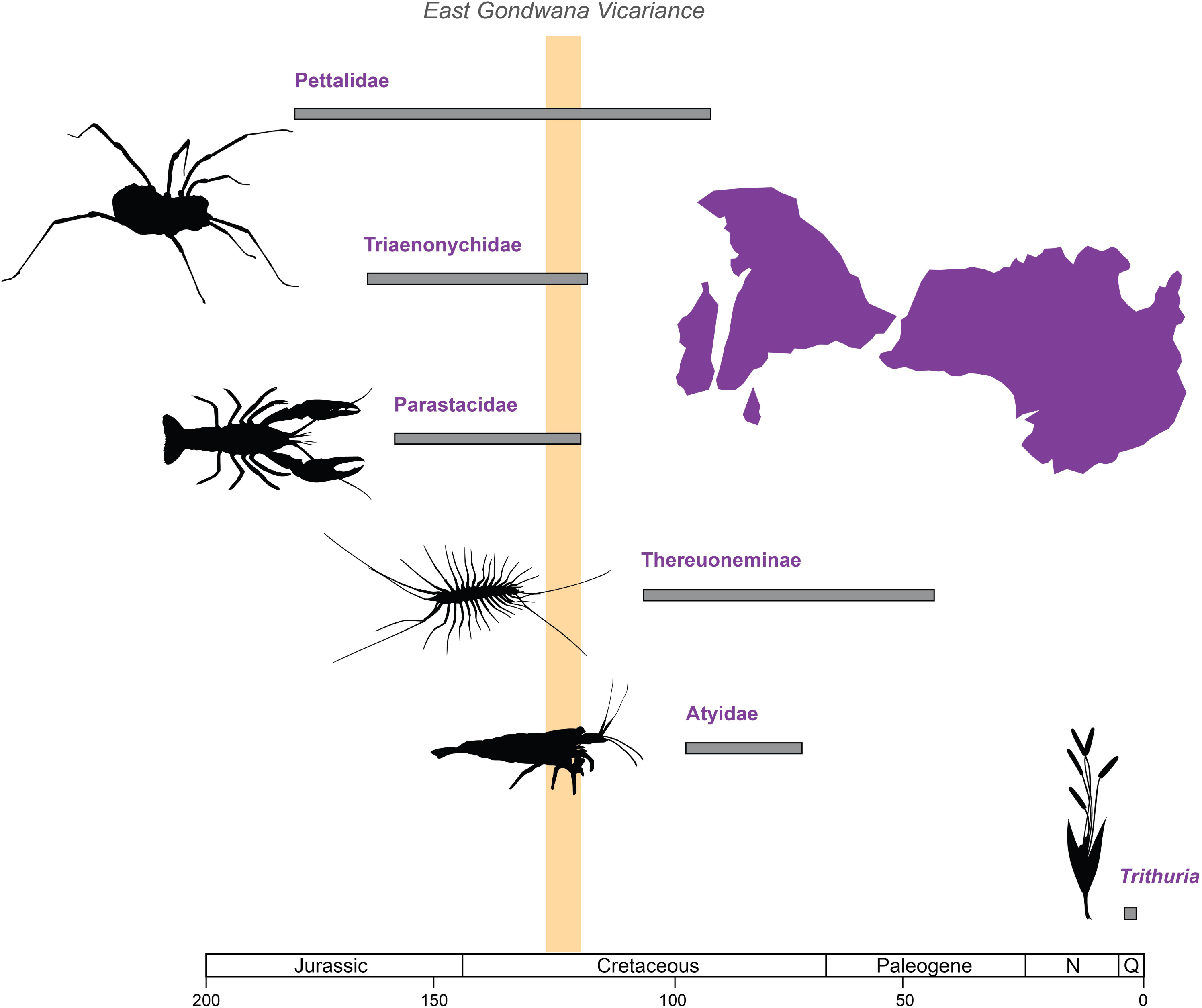
Examples of groups showing disjunct East Gondwana distribution. Gray bars correspond to credible intervals of divergence times between lineages with Peninsular Indian Plate/Malagasy and Australian affinities. Gold bar corresponds to the period of East Gondwana vicariance (120-130 mya). All silhouettes of taxa sourced from PhyloPic (Triaenonychidae: Jennifer Trimble, Parastacidae: Mattia Menchetti, *Scutigera*: Guillame Dera, Atyidae: Douglas Teles da Rosa, *Trithuria*: Michael Keesey) under Creative Commons License 4.0 (https://creativecommons.org/licenses/by/4.0/).

The PIP was contiguous with other East Gondwanan landmasses such as Madagascar, Seychelles, Australia and Antarctica until approximately 130 mya (Gibbons et al., 2013). At this time, PIP-Madagascar-Seychelles rifted from Australia-Antarctica. A few studies find evidence for the role of continental vicariance within East Gondwana in determining biodiversity patterns in some ancient, dispersal-limited taxa. For instance, Toon et al. (2010) show that the divergence of Madagascar crayfish (*Astacoides*) from Australian-NZ sister groups within the family Parastacidae occurred roughly between 121 and 160 mya. Baker, Boyer and Giribet (2020) find evidence for a vicariance event causing the divergence between the Australian *Austropurcellia* and Sri Lankan *Pettalus* of Pettalidae (a family of mite harvestmen with Gondwanan distribution) around 93-182 mya. Derkarabetian et al. (2021) also found that the Madagascar species *Ankaratrix illota* of the harvestman family Triaenonychidae diverged from Australian sisters between 119 and 166 mya. While the corresponding ancestral area reconstruction suggests an ancestral range of (Madagascar + Western Australia) coupled with extinction, East Gondwanan vicariance is not ruled out as a causal factor.

We find greater evidence in support of an ancient jump dispersal event between India and Australia around 44-107 mya. This is similar to results from Jurado-Rivera et al. (2017), where the divergence between the northwestern Australian *Stygiocaris* and the monotypic southwestern Malagasy *Typhlopatsa* cave shrimps is retrieved at 85.1 mya (CI: 97.9-72.7 mya), suggesting ancient trans-oceanic dispersal. On the other hand, for the aquatic plant family Hydatellaceae, which originated between 120.6-133.2 mya, Iles et al. (2014) show that the divergence between the sole Indian species *Trithuria konkanensis* and the northern Australian *T. lanterna* is a recent LDDE that occurred 0.76 (0.24-1.33) mya.

Our results are significant because the Cretaceous jump dispersal event from Peninsular India to Australia is likely to have driven the Australian-New Caledonian radiation in Scutigeromorpha, resulting in three genera that are monophyletic together- *Allothereua, Parascutigera and Pilbarascutigera*. While the date for this dispersal event overlaps with the timing of the Indo- Eurasian collision, it is unclear whether the dispersal route was transoceanic or on land. Sampling from Northeast India and Southeast Asia must be undertaken to solve this riddle; instances of an Australian/Pacific origin of taxa followed by dispersal into mainland Asia and India through Wallacea have been documented for leaf insects and a genus of butterflies, *Delias* (Bank et al., 2021; Braby & Pierce, 2007). Conversely, Xu et al. (2011) find evidence for the origin of *Ficus* groups Conosycea and Malvanthera in India, after which daughter lineages occupied SE Asia and Australia.

Biogeographic research on taxa with Indo-Australian distributions is an exciting possibility for future research. It would give us better insights into dispersal routes (land bridge, transoceanic, stepping-stone), direction of dispersal, and type of vicariance (continental, climatic) between the two Gondwanan landmasses through evolutionary time. In turn, this could help us identify common eco-evolutionary characteristics among taxa that allow them to show a biogeographic pattern that, so far in the literature, is rare indeed.

## Acknowledgements

We thank Abhishek Gopal, Dr Bharti Dharapuram, Dr Mihir Kulkarni, Payal Dash and Pragyadeep Roy for their insightful discussions on molecular techniques and analyses, assistance in fieldwork, and valuable feedback that significantly improved this manuscript. Additionally, we thank Bikash Sahoo for his help in the fieldwork. Field and lab work was supported by grants to JJ; The Wellcome Trust DBT India Alliance, Grant/Award Number: IA/I/20/1/504919; and The Council for Scientific and Industrial Research, Govt of India. We would also like to thank the state forest departments of Andhra Pradesh, Kerala, Karnataka, Maharashtra, Orissa, Tamil Nadu and Telangana for the research permits and their assistance during the fieldwork.

## Data availability statement

The DNA sequence data generated in this study are deposited in GenBank. The distribution data used for biogeography analyses are provided in Supplementary Materials S1 and S2.

## Biosketch

This study was part of Maya Manivannan’s Master’s dissertation at the Evolutionary Ecology Lab. Research at the Evo-Eco lab focuses on the macroecology and macroevolution of biodiversity in Tropical Asia, with particular interest in the systematics and biogeography of centipedes. See Evolutionary Ecology Lab, CSIR-CCMB Hyderabad, India, and Edgecombe at The Natural History Museum, United Kingdom, for more details.

## Author contributions

JJ and MM designed the study; JJ, NG, and MM conducted the fieldwork; MM generated the molecular data with the help of NG; MM and JJ analyzed the data, MM wrote the first draft with input from JJ and GE, and all the authors reviewed and edited the manuscript. JJ acquired the funding for this study.

## Appendices

### Appendix 1 DNA extraction and PCR

DNA was extracted from leg tissue using the Phenol Chloroform Isoamyl (PCI) method (Moore & Dowhan, 2002). Old samples that had become brittle from ethanol evaporation were digested in a lysis buffer formulated by (Santos et al., 2018). A lysis buffer containing 100 mM NaCl, 10 mM Tris-Cl, 1 mM disodium EDTA dihydrate, and 1% SDS was used for recently collected specimens. Two mitochondrial (Cytochrome *c* oxidase subunit I, 16S rDNA) and three nuclear (28S rDNA, 18S rDNA, Histone H3) markers were amplified in a 25 ul volume reaction in a Bio-Rad thermal cycler (T100) under the following conditions: initial denaturation at 94□ for 1.5 minutes, 35 cycles of denaturation at 95□ for 40s, annealing for 45s, extension at 72□ for 40s and final extension at 72□ for 6 minutes (see Table S3 for primer details). Amplification was verified with agarose gel electrophoresis.

### Appendix 2 Results of species delimitation among PIP scutigeromorphs

Within Clade 1 (n=9), a single species (*Thereuopoda* sp.; Species A) was recovered for the 16S dataset by ASAP, whereas PTP delimited four species. We lacked COI sequences for many individuals; therefore, we treated *Thereuopoda* sp. A is a single species distributed across the Western and Eastern Ghats. One of the largest clades among Indian Scutigeromorpha, Clade 2 (n=54), had three subclades. One subclade (n=5) was restricted to the southern parts of the Western Ghats, where two species based on COI datasets were recovered by ASAP and PTP. The 16S dataset recovered one species by ASAP and PTP and lacked the data for CCMB 1918, which was recovered as a distinct species in COI. Therefore, we treat them as two species in this subclade: Species B occurs in the southern Western Ghats at low and mid-elevation, and Species C is distributed at high elevation. Using the 16S dataset, ASAP recovered one of the largest subclades in Clade 2 (n=47) as a single species, whereas PTP recovered three distinct species. ASAP and PTP recovered five and six distinct clusters for the COI dataset. Given that ASAP recovered the subclade as a single distinct cluster, we treated it as one species, Species D, with distribution across Peninsular India. The third subclade of Clade 2 (n=2), consisting of two individuals from the northern Western Ghats, was recovered as a single unit by ASAP and PTP and as two distinct units by ASAP and PTP based on the 16S and COI datasets. Based on geographic proximity and monophyly in 16S and COI, we treated them as a single species in our analyses, Species E.

One of the clades of Thereuoneminae contained *Thereuopodina* sp. from Australia and an individual from the Andaman Islands (CCMB10), which were sister to an individual from the Eastern Ghats (CCMB4071). These individuals were recovered as three units from the 16S dataset by ASAP and PTP. As we lacked COI data, these were treated as three distinct species, Species F (CCMB4071), G (CCMB10) and *Thereuopodina* sp. Within Clade 3, two individuals from the central Western Ghats (CCMB1529 and CCMB442) were recovered as a single unit - Species H, by ASAP and PTP on 16S, although COI data were missing. Clade 4 consisted of four individuals. The COI dataset for three individuals recovered them as two distinct units by ASAP and PTP, Species J from the Eastern Ghats and K from the northern Western Ghats, respectively. Another individual from the central Western Ghats was recovered as a singleton (Species I) by ASAP and PTP from the 16S dataset but lacked COI data.

