## Supplementary figures for "A passage through India: the biotic ferry model supports the build-up of Indo-Australian biodiversity of an ancient soil arthropod clade"

Bayesian phylogeny of Scutigeromorpha based on COI, 16S, 28S, H3 and 18S makers. Posterior probability/UB2 bootstrapping supports are indicated at nodes. Names highlighted in bold indicate specimens from India. Photo of scutigeromorph by Nehal Gurung.

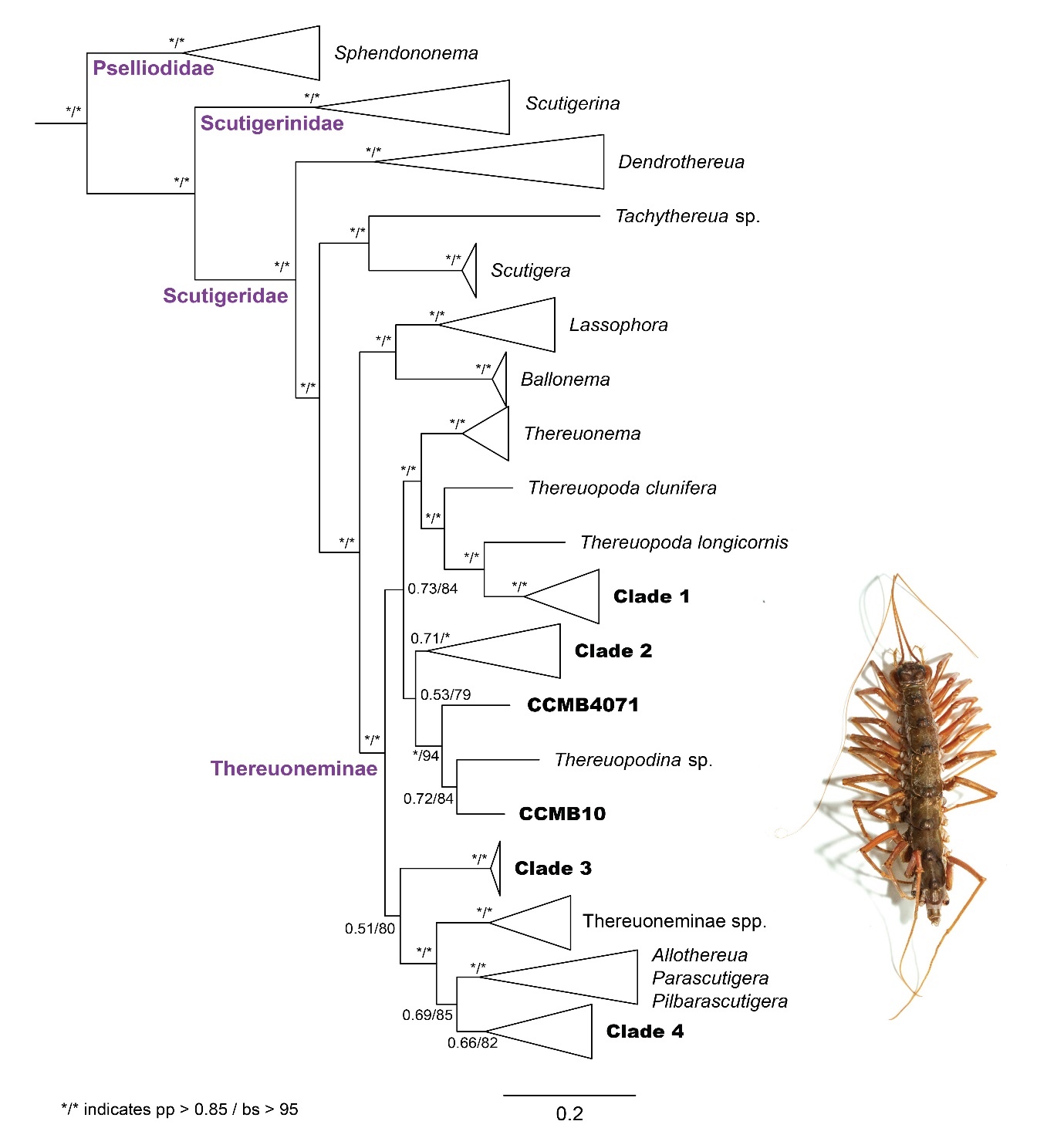
