## Supplementary figures for "A passage through India: the biotic ferry model supports the build-up of Indo-Australian biodiversity of an ancient soil arthropod clade"

### Table S1: Details of taxon sampling conducted in India.

**Note**: Clade identity from molecular phylogenetic analyses and species identity from species delimitation analyses for each specimen are also indicated. Columns labelled by marker show presence of molecular data (- indicates that data is absent); cells will be replaced with GenBank accession numbers.

| **Specimen** | **Clade** | **Species ID** | **Location** | **Longitude** | **Latitude** | **COI** | **16S** | **28S** | **H3** | **18S** |
| --- | --- | --- | --- | --- | --- | --- | --- | --- | --- | --- |
| CES07221 | 2 | D | Dandeli | 74.9603 | 15.27252 | **-** |  |  | **-** |  |
| CES1518 | 4 | I | Thenmala | 77.0651 | 8.9595 | **-** |  |  | **-** |  |
| CES1529 | 3 | H | Wayanad | 76.0832 | 11.704 | **-** |  |  |  |  |
| CCMB10 |  | G | Mount Harriet | 92.7335 | 11.7202 | **-** |  |  |  |  |
| CCMB100 | 1 | A | Lakhari Valley WLS | 84.32734 | 19.33728 | **-** |  |  |  |  |
| CCMB167 | 1 | A | Karlapat WLS | 84.02393 | 19.46843 | **-** |  |  |  |  |
| CCMB392 | 2 | D | Talacauvery, shola forest | 75.45979 | 12.39842 |  |  |  |  |  |
| CCMB442 | 3 | H | Talacauvery WLS | 75.45541 | 12.44458 |  |  | **-** |  |  |
| CCMB443 | 2 | D | Talacauvery WLS | 75.45543 | 12.44444 |  |  |  |  |  |
| CCMB466 | 2 | D | Talacauvery WLS | 75.45469 | 12.44381 |  |  |  |  |  |
| CCMB485 | 2 | D | Pushpagiri WLS | 75.70181 | 12.66079 |  |  |  |  |  |
| CCMB551 | 2 | D | Makutta Reserve Forest | 75.7608 | 12.08458 |  |  |  |  |  |
| CCMB552 | 2 | D | Makutta Reserve Forest | 75.7608 | 12.08458 |  |  |  |  |  |
| CCMB615 | 2 | D | Makutta Reserve Forest | 75.78353 | 12.12456 |  |  |  |  |  |
| CCMB616 | 2 | D | Makutta Reserve Forest | 75.78588 | 12.12418 |  | **-** |  |  |  |
| CCMB617 | 2 | D | Makutta Reserve Forest | 75.78588 | 12.12418 |  | **-** |  |  |  |
| CCMB618 | 2 | D | Makutta Reserve Forest | 75.78588 | 12.12418 |  |  |  |  |  |
| CCMB661 | 2 | D | Makutta Reserve Forest | 75.72576 | 12.07755 |  | **-** |  |  |  |
| CCMB664 | 2 | D | Makutta Reserve Forest | 75.72576 | 12.07755 |  |  |  |  |  |
| CCMB701 | 2 | D | Makutta Reserve Forest | 75.71429 | 12.14618 |  | **-** |  |  |  |
| CCMB702 | 2 | D | Makutta Reserve Forest | 75.71429 | 12.14618 |  |  |  |  |  |
| CCMB703 | 2 | D | Makutta Reserve Forest | 75.70841 | 12.13926 |  |  |  |  |  |
| CCMB704 | 2 | D | Makutta Reserve Forest | 75.70841 | 12.13926 |  |  |  |  |  |
| CCMB705 | 2 | D | Makutta Reserve Forest | 75.71429 | 12.14618 |  |  |  |  |  |
| CCMB717 | 2 | D | Brahmagiri WLS, Abayalu | 75.87815 | 11.98819 |  |  |  |  |  |
| CCMB733 | 2 | D | Brahmagiri WLS, Abayalu | 75.87586 | 11.98536 |  |  |  | **-** |  |
| CCMB747 | 2 | D | Anshi WLS | 74.38542 | 14.98568 |  |  |  |  |  |
| CCMB749 | 2 | D | Anshi WLS | 74.38542 | 14.98568 |  |  |  |  |  |
| CCMB765 | 4 | K | Van Kusavade | 73.9199 | 17.5652 |  | **-** |  |  |  |
| CCMB798 | 1 | A | Ranebennur WLS | 75.66452 | 14.649282 | **-** |  |  |  |  |
| CCMB880 | 2 | D | Anshi WLS, Maingini Trek | 74.39169 | 15.01573 |  |  |  |  |  |
| CCMB897 | 2 | D | Anshi WLS, Badpoli | 74.36226 | 14.99488 |  |  |  |  |  |
| CCMB898 | 2 | D | Anshi WLS, Badpoli | 74.36226 | 14.99488 |  |  |  |  |  |
| CCMB899 | 2 | D | Anshi WLS, Badpoli | 74.35898 | 14.9949 |  |  |  |  |  |
| CCMB900 | 2 | D | Anshi WLS, Badpoli | 74.35898 | 14.9949 |  | **-** |  |  |  |
| CCMB939 | 2 | D | Castlerock, Kuveshi | 74.31981 | 15.3211 |  |  |  |  |  |
| CCMB973 | 2 | D | Hejani | 74.75882 | 14.25725 |  |  |  |  |  |
| CCMB1003 | 2 | D | Kathlekan | 74.74619 | 14.27571 |  |  |  |  |  |
| CCMB1069 | 2 | D | Someshwara WLS, Badimatta | 75.06815 | 13.45911 |  |  |  |  |  |
| CCMB1098 | 2 | D | Agumbe | 75.98723 | 13.51449 |  |  |  | **-** |  |
| CCMB1125 | 2 | D | Mookambika WLS, Aanejary | 74.80003 | 13.83274 |  |  |  |  |  |
| CCMB1126 | 2 | D | Mookambika WLS, Aanejary | 74.80003 | 13.83274 |  |  |  |  |  |
| CCMB1202 | 1 | A | Gundlabramhmeswaram WLS | 78.81849 | 15.453 | **-** |  |  |  |  |
| CCMB1231 | 1 | A | Gundlabramhmeswaram WLS | 78.81849 | 15.453 | **-** |  |  |  |  |
| CCMB1347 | 2 | D | Silent Valley National Park | 76.45467 | 11.09576 |  |  | **-** |  |  |
| CCMB1353 | 2 | D | Silent Valley National Park | 76.45467 | 11.09576 |  |  |  |  |  |
| CCMB1404 | 2 | D | Silent Valley National Park | 76.45467 | 11.09576 |  |  |  |  |  |
| CCMB1474 | 2 | D | Siruvani | 76.65784 | 10.9715 |  |  |  | **-** |  |
| CCMB1534 | 2 | D | Wayanad WLS | 76.34615 | 11.70722 |  |  |  |  |  |
| CCMB1591 | 2 | D | Wayanad WLS | 76.09776 | 11.91699 |  | **-** |  |  |  |
| CCMB1644 | 2 | D | Aralam WLS | 75.81257 | 11.93311 |  |  |  |  |  |
| CCMB1663 | 2 | D | Aralam WLS | 75.88126 | 11.94627 |  |  |  |  |  |
| CCMB1664 | 2 | D | Aralam WLS | 75.8501 | 11.93731 |  |  |  |  |  |
| CCMB1673 | 2 | D | Aralam WLS | 75.88126 | 11.94627 |  |  |  |  |  |
| CCMB1918 | 2 | C | Eravikulam National Park | 77.080320 | 10.156890 |  | **-** |  |  |  |
| CCMB2029 | 2 | B | Parambikulam Tiger Reserve | 76.623180 | 10.384560 |  |  |  |  |  |
| CCMB2030 | 2 | D | Parambikulam Tiger Reserve | 76.625490 | 10.381240 |  | **-** |  |  |  |
| CCMB2031 | 2 | B | Parambikulam Tiger Reserve | 76.625490 | 10.381240 |  |  |  |  |  |
| CCMB2032 | 2 | D | Parambikulam Tiger Reserve | 76.623180 | 10.384560 |  | **-** |  |  |  |
| CCMB2073 | 2 | B | Parambikulam Tiger Reserve | 76.623180 | 10.384560 | **-** |  |  | **-** |  |
| CCMB2082 | 2 | D | Parambikulam Tiger Reserve | 76.783270 | 10.420280 |  |  |  | **-** |  |
| CCMB2126 | 2 | B | Parambikulam Tiger Reserve | 76.687540 | 10.304760 | **-** |  |  |  |  |
| CCMB2227 | 2 | E | Radhanagari WLS | 73.95871 | 16.35068 |  |  |  |  |  |
| CCMB2228 | 1 | A | Radhanagari WLS | 73.95537 | 16.34891 | **-** |  |  |  |  |
| CCMB2229 | 2 | E | Radhanagari WLS | 73.95537 | 16.34891 |  |  |  |  |  |
| CCMB2230 | 1 | A | Radhanagari WLS | 73.94569 | 16.34603 | **-** |  |  |  |  |
| CCMB2298 | 4 | J | Amrabad Tiger Reserve | 78.70561 | 16.25707 |  | **-** |  | **-** |  |
| CCMB2299 | 4 | J | Amrabad Tiger Reserve | 78.70561 | 16.25707 |  | **-** |  | **-** |  |
| CCMB4070 | 1 | A | Odisha Eastern Ghats | 84.2039 | 20.442 | **-** |  |  |  |  |
| CCMB4071 |  | F | Odisha Eastern Ghats | 84.2039 | 20.442 | **-** |  |  |  |  |
| CCMB4072 | 1 | A | NISER Campus, Jatni | 85.6861 | 20.17201 |  |  |  |  |  |

### Table S2: GenBank sequences for phylogenetic analysis.

**Note**: All data has been published previously by Giribet and Edgecombe (2013).

| **Family** | **Species** | **Voucher no** | **Location** | **LON** | **LAT** | **COI** | **16S** | **28S** | **18S** | **H3** |
| --- | --- | --- | --- | --- | --- | --- | --- | --- | --- | --- |
| Outgroup | *Proteroiulus fuscus* | DNA100171 |  |  |  | AF370842 | AF370866 | AF370804 | AF173236 |  |
|  | *Lithobius variegatus* |  |  |  |  | AF334311 | AY084071 | AF000780 | AF000773 |  |
|  | *Anopsobius neozelanicus* | DNA101035 |  |  |  | DQ222165 | AF334337 | DQ222132 | AF173248 | DQ222179 |
|  | *Paralamyctes validus* | DNA1 00297 |  |  |  | AF334330 | AF334358 | AF173278 | AF334289 | DQ222180 |
|  | *Craterostigmus tasmanius* | DNA100280 |  |  |  | AF370835 | AF370860 | DQ222133 | AF000774 | AF110850 |
|  | *Scolopendra viridis* | DNA100675 |  |  |  | DQ201431 | DQ201425 | DQ222134 | DQ201419 | DQ222181 |
| Scutigeromorpha |  |  |  |  |  |  |  |  |  |  |
| Pselliodidae | *Sphendononema guildingii* | DNA101161 | Brazil | -38.831 | -19.561 |  |  | DQ222138 | DQ222121 | DQ222185 |
|  | *Sphendononema guildingii* | DNA101630 | Brazil | -39.1023 | -16.589 | DQ222168 | DQ222154 | DQ222139 | DQ222122 | DQ222186 |
|  | *Sphendononema rugosa* | DNA104623 | Cameroon | 12.867 | 5.9694 | HQ591450 | HQ591448 | HQ591447 | HQ591445 | HQ591451 |
| Scutigerinidae | *Scutigerina hova* | DNA101592 | Madagascar | 47.2869 | -18.225 |  | DQ222153 | DQ222137 | DQ222120 | DQ222184 |
|  | *Scutigerina malagassa* | DNA101591 | Madagascar | 47.2819 | -18.171 | DQ222167 | DQ222152 | KF218758 | DQ222119 | DQ222183 |
|  | *Scutigerina weberi* | DNA100455 | Swaziland | 31.1189 | -25.997 | AY288741 | AY288717 |  | AY288689 | AY428835 |
|  | *Scutigerina weberi* | DNA106731 | South Africa | 22.4175 | -30.596 | KF218788 | KF218776 | KF218759 | KF218741 | KF218801 |
|  | *Scutigerina weberi* | DNA106732 | South Africa | 22.4175 | -30.596 |  | KF218777 | KF218760 | KF218742 | KF218802 |
|  | *Scutigerina weberi* | DNA106733 | South Africa | 22.4175 | -30.596 | KF218789 |  | KF218761 | KF218743 |  |
|  | *Scutigerina cf. weberi* | DNA101590 | Madagascar | 47.4225 | -21.25 | DQ222166 | DQ222151 |  | DQ222118 | DQ222182 |
| Scutigeridae, Scutigerinae | *Dendrothereua homa* | DNA102576 | USA | -110.665 | 32.033 | FJ660818 | FJ660786 | FJ660746 | FJ660705 | FJ660842 |
|  | *Dendrothereua nubila* | DNA101791 | Costa Rica | -83.9594 | 10.007 | FJ660817 | FJ660785 | FJ660745 | FJ660704 | FJ660841 |
|  | *Dendrothereua* sp. | DNA106729 | Mexico | -101.726 | 23.795 | KF218790 | KF218778 |  | KF218744 | KF218803 |
|  | *Dendrothereua* sp. | DNA106730 | Mexico | -101.726 | 23.795 | KF218791 | KF218779 | KF218767 | KF218745 |  |
|  | *Dendrothereua* sp. | DNA106854 | Mexico | -101.726 | 23.795 | KF218792 |  | KF218768 | KF218746 |  |
|  | *Dendrothereua* sp. | DNA107050 | Dominican Rep | -69.9467 | 18.929 |  | KF218780 | KF218769 | KF218747 | KF218804 |
|  | *Dendrothereua* sp. | IZ89420 | Guatemala | -90.3377 | 15.6557 | KF218794 | KF218781 | KF218770 | KF218748 |  |
|  | *Dendrothereua* sp. | IZ125268 | Nicaragua | -84.5937 | 12.9805 | KF218793 |  | KF218772 | KF218749 |  |
|  | *Scutigera coleoptrata* | DNA100198 | USA | -71.1134 | 42.3792 |  | DQ222155 | DQ222140 | DQ222123 | DQ222187 |
|  | *Scutigera coleoptrata* | DNA100259 | South Africa | 18.4618 | -33.957 | DQ222170 | DQ222156 | EF199983 | DQ222124 |  |
|  | *Scutigera coleoptrata* | DNA102327 | Turkey | 32.7599 | 36.1167 |  | FJ660788 | FJ660748 | FJ660707 | FJ660844 |
|  | *Scutigera coleoptrata* | DNA102329 | Georgia | 42.5533 | 42.1978 |  | FJ660789 | FJ660750 | FJ660709 | FJ660846 |
|  | *Scutigera coleoptrata* | DNA102367 | Bulgaria | 26.5009 | 42.4845 |  | FJ660790 | FJ660751 | FJ660710 | FJ660847 |
|  | *Scutigera coleoptrata* | DNA102368 | Bulgaria | 26.5009 | 42.4845 | FJ660820 | FJ660791 | FJ660752 | FJ660711 | FJ660848 |
|  | *Scutigera coleoptrata* | DNA102577 | Elba | 10.1466 | 42.7343 |  | FJ660792 | FJ660753 | FJ660712 | FJ660849 |
|  | *Scutigera coleoptrata* | DNA102578 | France | 0.2678 | 44.7772 |  | FJ660793 | FJ660754 | FJ660713 | FJ660850 |
| Scutigeridae, Incertae cedis | *Ballonema* sp. | DNA106728 | PNG | 144.1873 | -5.8768 | KF218796 | KF218782 | KF218762 | KF218751 | KF218805 |
|  | *Ballonema* sp. | DNA106902 | PNG | 144.1873 | -5.8768 | KF218795 | KF218783 | KF218763 | KF218752 | KF218806 |
|  | *Ballonema* sp. | DNA106903 | PNG | 144.1873 | -5.8768 | KF218797 | KF218784 | KF218764 | KF218753 | KF218807 |
|  | *Lassophora nossibei* | DNA102102 | Madagascar | 46.8067 | -16.323 | FJ660821 | FJ660794 | FJ660755 | FJ660715 | FJ660851 |
|  | *Lassophora* sp. | DNA106727 | Mozambique | 35.0315 | -17.166 | KF218798 | KF218785 | KF218773 | KF218754 | KF218808 |
|  | *Tachythereua* sp. | DNA102575 | Senegal | -16.9564 | 14.0873 |  | FJ660795 | FJ660756 | FJ660716 | FJ660852 |
| Scutigeridae, Thereuoneminae | *Allothereua bidenticulata* | DNA101589 | Australia | 151.065 | -33.98 | FJ660822 | FJ660796 | FJ660757 | FJ660717 |  |
|  | *Allothereua linderi* | DNA101463 | Australia | 146.0167 | -35.98 | DQ222174 | DQ222160 | DQ222147 | DQ222128 | DQ222195 |
|  | *Allothereua linderi* | DNA101979 | Australia | 146.35 | -38.145 |  | FJ660797 | FJ660758 | FJ660719 | FJ660853 |
|  | *Allothereua maculata* | DNA101982 | Australia | 116.0417 | -34.54 | FJ660823 | FJ660798 | FJ660759 | FJ660720 | FJ660854 |
|  | *Allothereua maculata* | DNA101983 | Australia | 116.0417 | -34.54 | FJ660824 | FJ660799 | FJ660760 | FJ660721 | FJ660855 |
|  | *Allothereua maculata* | DNA101986 | Australia | 115.7244 | -33.47 | FJ660825 | FJ660800 | FJ660761 | FJ660722 | FJ660856 |
|  | *Allothereua maculata* | DNA101987 | Australia | 115.37 | -34.251 | FJ660826 | FJ660801 | FJ660762 | FJ660723 |  |
|  | *Allothereua maculata* | DNA101988 | Australia | 115.9667 | -33.083 | FJ660827 | FJ660802 | FJ660763 | FJ660724 | FJ660857 |
|  | *Allothereua serrulata* | DNA100262 | Australia | 152.6097 | -26.667 | DQ222175 | DQ222161 | DQ222148 | DQ222129 | DQ222197 |
|  | *Allothereua serrulata* | DNA101045 | Australia | 145.0239 | -22.617 | DQ222176 | DQ222162 | FJ660764 | DQ222130 | DQ222198 |
|  | *Parascutigera festiva* | DNA100635 | New Caledonia | 166.15 | -21.75 | DQ222177 |  | FJ660765 | AY288688 | DQ222199 |
|  | *Parascutigera festiva* | DNA102584 | New Caledonia | 164.9119 | -20.952 | FJ660828 | FJ660803 | FJ660766 | FJ660725 |  |
|  | *Parascutigera guttata* | DNA101971 | Australia | 145.1833 | -16.817 |  | FJ660805 | FJ660768 | FJ660727 |  |
|  | *Parascutigera guttata* | DNA101973 | Australia | 145.2667 | -16.567 |  | FJ660806 | FJ660769 | FJ660729 |  |
|  | *Parascutigera guttata* | DNA102317 | Australia | 145.7022 | -17.314 | FJ660829 | FJ660804 | FJ660767 | FJ660726 | FJ660859 |
|  | *Parascutigera latericia* | DNA101046 | New Caledonia | 164.5167 | -20.4 | DQ222178 | DQ222164 | DQ222150 | DQ222131 | DQ222200 |
|  | *Parascutigera latericia* | DNA102123 | New Caledonia | 165.3167 | -21.183 | FJ660830 | FJ660807 |  | FJ660730 | FJ660860 |
|  | *Parascutigera latericia* | DNA102124 | New Caledonia | 166.6833 | -22.217 | FJ660831 |  | FJ660771 | FJ660731 |  |
|  | *Parascutigera nubila* | DNA103553 | New Caledonia | 166.65 | -22.117 | FJ660832 | FJ660808 | FJ660772 | FJ660732 |  |
|  | *Parascutigera* sp. QLD 1 | DNA101974 | Australia | 148.85 | -21.767 | FJ660834 | FJ660809 | FJ660775 | FJ660735 | FJ660861 |
|  | *Parascutigera* sp. QLD 2 | DNA101972 | Australia | 147.8733 | -22.477 |  | FJ660810 | FJ660776 | FJ660736 | FJ660862 |
|  | *Parascutigera* sp. QLD 3 | DNA101977 | Australia | 145.5723 | -17.288 | FJ660835 | FJ660811 | FJ660777 | FJ660737 | FJ660863 |
|  | *Parascutigera* sp. QLD 3 | DNA101978 | Australia | 145.4994 | -17.333 | FJ660836 | FJ660812 | FJ660778 | FJ660738 | FJ660864 |
|  | *Parascutigera cf. sphinx* | DNA101980 | Australia | 116.0767 | -32.763 | FJ660838 | FJ660813 | FJ660780 | FJ660740 | FJ660866 |
|  | *Parascutigera cf. sphinx* | DNA101981 | Australia | 116.0767 | -32.763 | FJ660839 | FJ660814 | FJ660781 | FJ660741 | FJ660867 |
|  | *Parascutigera cf. sphinx* | DNA101985 | Australia | 116.0958 | -32.404 | FJ660837 |  | FJ660779 | FJ660739 | FJ660865 |
|  | *PIPlbarascutigera incola* | DNA101997 | Australia | 119.3811 | -22.298 |  | FJ660815 | FJ660783 | FJ660742 |  |
|  | *Thereuopoda clunifera* | DNA100260 | Japan | 137.896 | 35.9396 | DQ222171 | AY288716 | DQ222142 | AF173239 | DQ222190 |
|  | *Thereuopoda longicornis* | DNA101461 | Thailand | 98.8297 | 19.3203 | DQ222172 | DQ222157 | DQ222143 | DQ222125 | DQ222191 |
|  | *Thereuonema tuberculata* | DNA101632 | Japan | 134.2145 | 35.3665 | DQ222173 | DQ222158 | DQ222145 | DQ222126 | DQ222193 |
|  | *Thereuonema turkestana* | DNA101090 | Kazakhstan | 77.0667 | 43.95 | FJ660840 | FJ660816 | FJ660784 | FJ660743 |  |
|  | *Thereuonema turkestana* | DNA101091 | Uzbekistan | 66.8729 | 39.5667 | DQ201427 | DQ201423 | DQ222144 | DQ201417 | DQ222192 |
|  | Thereuoneminae sp. | IZ103188 | Guam | 144.7831 | 13.4502 | KF218799 | KF218786 | KF218774 | KF218756 | KF218809 |
|  | Thereuoneminae sp. | IZ103200 | Micronesia | 151.0102 | 11.8389 | KF218800 | KF218787 | KF218775 | KF218757 | KF218810 |
|  | *Thereuopodina* | DNA101462 | Australia | 144.498 | -17.183 |  | DQ222159 | DQ222146 | DQ222127 | DQ222194 |

### Table S3: Primer details and conditions

| **Marker** | **Amplicon size (bp)** | **Primer** | **Primer reference** | **Sequence (5’- 3’)** | **Annealing temperature (℃)** |
| --- | --- | --- | --- | --- | --- |
| COI | 656 | LCOI490 | Folmer et al. (1994) | GGTCAACAAATCATAAAGATATTGG | 46 |
|  |  | HCO2198 |  | TAAACTTCAGGGTGACCAAAAAATCA |  |
| 16S rRNA | 531 | 16Sar | Xiong and Kocher (1991) | CGCCTGTTTATCAAAAACAT | 49 |
|  |  | 16Sb |  | CTCCGGTTTGAACTCAGATCA |  |
| 28S rRNA | 486 | 28Sa | Whiting et al. (1997) | GACCCGTCTTGAAACACGGA | 52 |
|  |  | 28Sb |  | TCGGAAGGAACCAGCTAC |  |
| 18S rRNA | 327 | 18S_2F | Machida and Knowlton (2012) | AACTTAAAGRAATTGACGGA | 56 |
|  |  | 18S_4R |  | CKRAGGGCATYACWGACCTGTTAT |  |
| H3 | 354 | H3aF | Colgan et al. (1998) | ATGGCTCGTACCAAGCAGACVGC | 54 |
|  |  | H3aR |  | ATATCCTTRGGCATRATRGTGAC |  |
|  | **2354** |  |  |  |  |

### Table S4: Fossil calibrations for divergence time estimation

**Note:** All fossil calibrations noted below were obtained from Wolfe et al. (2016)

| **Sl.** | **Fossil** | **Phylogenetic position** | **Age interval (mya)** |
| --- | --- | --- | --- |
| 1 | *Cowiedesmus eroticopodus* | Root | 424.7 - 636.1 |
| 2 | *Crussolum* | Crown Chilopoda | 416 - 636.1 |
| 3 | *Devonobius delta* | Crown Pleurostigmophora | 382.7 - 521 |
| 4 | *Mazoscolopendra richardsoni* | Crown Amalphighiata | 306.9 - 521 |
| 5 | *Fulmenocursor tenax* | Crown Scutigeromorpha | 112.6 - 521 |

### Table S5: BioGeoBEARS model setup

**Time points**: 150 mya (rifting between East and West Gondwana), 120 mya (rifting between PIP and the rest of East Gondwana), 100 mya (rifting between the Neotropics and Africa), 40 mya (collision between PIP and the Palearctic, the emergence of New Caledonia and Guam) and 15 mya (emergence of the Andaman and Nicobar Islands)

**Area codes**: Africa (Afr), Australia (Aus), Madagascar (MDG), Pacific Islands (Pac Isl), Nearctic (Nea), Neotropics (Neo), New Caledonia (NC), Palearctic (Pal), Peninsular Indian Plate (PIP), SE Asia (SE), Andaman Islands (AN). Refer to text for geographic delimitations.

**Keys**:

| **Condition** | **Dispersal multiplier** |
| --- | --- |
| Same area/ Adjacent areas | 1 |
| Separated by small water barrier | 0.75 |
| Separated by another area | 0.5 |
| Separated by large water barrier OR  by a small water barrier + land area OR  large terrestrial barrier | 0.25 |
| Area does not exist | 0 |

**Dispersal matrix:**

| **360 mya (tree root)** | | | | | | | | | | | |
| --- | --- | --- | --- | --- | --- | --- | --- | --- | --- | --- | --- |
|  | Afr | Aus | MDG | Pac Isl | Nea | Neo | NC | Pal | PIP | SE | AN |
| Afr | 1 | 0.5 | 1 | 0 | 0.25 | 1 | 0 | 0.5 | 1 | 1 | 0 |
| Aus | 0.5 | 1 | 0.5 | 0 | 0.25 | 0.25 | 0 | 0.25 | 1 | 1 | 0 |
| MDG | 1 | 0.5 | 1 | 0 | 0.25 | 0.5 | 0 | 0.25 | 1 | 0.5 | 0 |
| Pac Isl | 0 | 0 | 0 | 0 | 0 | 0 | 0 | 0 | 0 | 0 | 0 |
| Nea | 0.25 | 0.25 | 0.25 | 0 | 1 | 0.75 | 0 | 1 | 0.25 | 0.25 | 0 |
| Neot | 1 | 0.25 | 0.5 | 0 | 0.75 | 1 | 0 | 0.75 | 0.25 | 0.5 | 0 |
| NC | 0 | 0 | 0 | 0 | 0 | 0 | 0 | 0 | 0 | 0 | 0 |
| Pal | 0.5 | 0.25 | 0.25 | 0 | 1 | 0.75 | 0 | 1 | 0.25 | 0.25 | 0 |
| PIP | 1 | 1 | 1 | 0 | 0.25 | 0.25 | 0 | 0.25 | 1 | 1 | 0 |
| SE | 1 | 1 | 0.5 | 0 | 0.25 | 0.5 | 0 | 0.25 | 1 | 1 | 0 |
| AN | 0 | 0 | 0 | 0 | 0 | 0 | 0 | 0 | 0 | 0 | 0 |
| **150 mya (East and West Gondwana rift)** | | | | | | | | | | | |
|  | Afr | Aus | MDG | Pac Isl | Nea | Neo | NC | Pal | PIP | SE | AN |
| Afr | 1 | 0.25 | 0.75 | 0 | 0.5 | 1 | 0 | 1 | 0.75 | 0.5 | 0 |
| Aus | 0.25 | 1 | 0.5 | 0 | 0.25 | 0.25 | 0 | 0.25 | 1 | 0.75 | 0 |
| MDG | 0.75 | 0.5 | 1 | 0 | 0.25 | 0.5 | 0 | 0.25 | 1 | 0.25 | 0 |
| Pac Isl | 0 | 0 | 0 | 0 | 0 | 0 | 0 | 0 | 0 | 0 | 0 |
| Nea | 0.5 | 0.25 | 0.25 | 0 | 1 | 1 | 0 | 1 | 0.25 | 0.5 | 0 |
| Neot | 1 | 0.25 | 0.25 | 0 | 1 | 1 | 0 | 0.5 | 0.5 | 0.25 | 0 |
| NC | 0 | 0 | 0 | 0 | 0 | 0 | 0 | 0 | 0 | 0 | 0 |
| Pal | 1 | 0.25 | 0.25 | 0 | 1 | 0.5 | 0 | 1 | 0.25 | 1 | 0 |
| PIP | 0.75 | 1 | 1 | 0 | 0.25 | 0.5 | 0 | 0.25 | 1 | 0.25 | 0 |
| SE | 0.5 | 1 | 0.25 | 0 | 0.5 | 0.25 | 0 | 1 | 0.5 | 1 | 0 |
| AN | 0 | 0 | 0 | 0 | 0 | 0 | 0 | 0 | 0 | 0 | 0 |
| **120 mya (PIP rifts from the rest of East Gondwana)** | | | | | | | | | | | |
|  | Afr | Aus | MDG | Pac Isl | Nea | Neo | NC | Pal | PIP | SE | AN |
| Afr | 1 | 0.25 | 0.75 | 0 | 0.5 | 1 | 0 | 1 | 0.75 | 0.25 | 0 |
| Aus | 0.25 | 1 | 0.5 | 0 | 0.25 | 0.25 | 0 | 0.25 | 0.75 | 0.25 | 0 |
| MDG | 0.75 | 0.5 | 1 | 0 | 0.25 | 0.5 | 0 | 0.25 | 1 | 0.25 | 0 |
| Pac Isl | 0 | 0 | 0 | 0 | 0 | 0 | 0 | 0 | 0 | 0 | 0 |
| Nea | 0.5 | 0.25 | 0.25 | 0 | 1 | 1 | 0 | 1 | 0.25 | 0.25 | 0 |
| Neot | 1 | 0.25 | 0.5 | 0 | 1 | 1 | 0 | 0.5 | 0.25 | 0.25 | 0 |
| NC | 0 | 0 | 0 | 0 | 0 | 0 | 0 | 0 | 0 | 0 | 0 |
| Pal | 1 | 0.25 | 0.25 | 0 | 1 | 0.5 | 0 | 1 | 0.25 | 1 | 0 |
| PIP | 0.75 | 0.75 | 1 | 0 | 0.25 | 0.25 | 0 | 0.25 | 1 | 0.25 | 0 |
| SE | 0.5 | 0.25 | 0.25 | 0 | 0.25 | 0.25 | 0 | 1 | 0.25 | 1 | 0 |
| AN | 0 | 0 | 0 | 0 | 0 | 0 | 0 | 0 | 0 | 0 | 0 |
| **100 mya (Africa and South America rift)** | | | | | | | | | | | |
|  | Afr | Aus | MDG | Pac Isl | Nea | Neo | NC | Pal | PIP | SE | AN |
| Afr | 1 | 0.25 | 0.75 | 0 | 0.5 | 0.75 | 0 | 1 | 0.25 | 0.25 | 0 |
| Aus | 0.25 | 1 | 0.25 | 0 | 0.25 | 0.25 | 0 | 0.25 | 0.25 | 0.25 | 0 |
| MDG | 0.75 | 0.25 | 1 | 0 | 0.25 | 0.25 | 0 | 0.25 | 1 | 0.25 | 0 |
| Pac Isl | 0 | 0 | 0 | 0 | 0 | 0 | 0 | 0 | 0 | 0 | 0 |
| Nea | 0.5 | 0.25 | 0.25 | 0 | 1 | 1 | 0 | 1 | 0.25 | 0.5 | 0 |
| Neot | 0.75 | 0.25 | 0.25 | 0 | 1 | 1 | 0 | 0.5 | 0.25 | 0.25 | 0 |
| NC | 0 | 0 | 0 | 0 | 0 | 0 | 0 | 0 | 0 | 0 | 0 |
| Pal | 1 | 0.25 | 0.25 | 0 | 1 | 0.5 | 0 | 1 | 0.25 | 1 | 0 |
| PIP | 0.25 | 0.25 | 1 | 0 | 0.25 | 0.25 | 0 | 0.25 | 1 | 0.25 | 0 |
| SE | 0.5 | 0.25 | 0.25 | 0 | 0.5 | 0.25 | 0 | 1 | 0.25 | 1 | 0 |
| AN | 0 | 0 | 0 | 0 | 0 | 0 | 0 | 0 | 0 | 0 | 0 |
| **40 mya (Collision between PIP and Eurasia; emergence of NC and Guam)** | | | | | | | | | | | |
|  | Afr | Aus | MDG | Pac Isl | Nea | Neo | NC | Pal | PIP | SE | AN |
| Afr | 1 | 0.25 | 0.75 | 0.25 | 0.25 | 0.25 | 0.25 | 1 | 0.5 | 0.5 | 0 |
| Aus | 0.25 | 1 | 0.25 | 0.25 | 0.25 | 0.25 | 0.75 | 0.25 | 0.25 | 0.25 | 0 |
| MDG | 0.75 | 0.25 | 1 | 0.25 | 0.25 | 0.25 | 0.25 | 0.25 | 0.25 | 0.25 | 0 |
| Pac Isl | 0.25 | 0.25 | 0.25 | 1 | 0.25 | 0.25 | 0.25 | 0.25 | 0.25 | 0.25 | 0 |
| Nea | 0.25 | 0.25 | 0.25 | 0.25 | 1 | 1 | 0.25 | 0.75 | 0.25 | 0.5 | 0 |
| Neot | 0.25 | 0.25 | 0.25 | 0.25 | 1 | 1 | 0.25 | 0.25 | 0.25 | 0.25 | 0 |
| NC | 0.25 | 0.75 | 0.25 | 0.25 | 0.25 | 0.25 | 1 | 0.25 | 0.25 | 0.25 | 0 |
| Pal | 1 | 0.25 | 0.25 | 0.25 | 0.75 | 0.25 | 0.25 | 1 | 1 | 1 | 0 |
| PIP | 0.75 | 0.25 | 0.25 | 0.25 | 0.25 | 0.25 | 0.25 | 1 | 1 | 0.75 | 0 |
| SE | 0.5 | 0.25 | 0.25 | 0.25 | 0.5 | 0.25 | 0.25 | 1 | 0.75 | 1 | 0 |
| AN | 0 | 0 | 0 | 0 | 0 | 0 | 0 | 0 | 0 | 0 | 0 |
| **15 mya (Emergence of the Andaman and Nicobar Islands)** | | | | | | | | | | | |
|  | Afr | Aus | MDG | Pac Isl | Nea | Neo | NC | Pal | PIP | SE | AN |
| Afr | 1 | 0.25 | 0.75 | 0.25 | 0.25 | 0.25 | 0.25 | 1 | 0.5 | 0.5 | 0.25 |
| Aus | 0.25 | 1 | 0.25 | 0.25 | 0.25 | 0.25 | 0.75 | 0.25 | 0.5 | 0.75 | 0.5 |
| MDG | 0.75 | 0.25 | 1 | 0.25 | 0.25 | 0.25 | 0.25 | 0.25 | 0.25 | 0.25 | 0.25 |
| Pac Isl | 0.25 | 0.25 | 0.25 | 1 | 0.25 | 0.25 | 0.25 | 0.25 | 0.25 | 0.25 | 0.25 |
| Nea | 0.25 | 0.25 | 0.25 | 0.25 | 1 | 1 | 0.25 | 0.25 | 0.25 | 0.25 | 0.25 |
| Neot | 0.25 | 0.25 | 0.25 | 0.25 | 1 | 1 | 0.25 | 0.25 | 0.25 | 0.25 | 0.25 |
| NC | 0.25 | 0.75 | 0.25 | 0.25 | 0.25 | 0.25 | 1 | 0.25 | 0.25 | 0.25 | 0.25 |
| Pal | 1 | 0.25 | 0.25 | 0.25 | 0.25 | 0.25 | 0.25 | 1 | 1 | 1 | 0.5 |
| PIP | 0.75 | 0.5 | 0.25 | 0.25 | 0.25 | 0.25 | 0.25 | 1 | 1 | 1 | 0.5 |
| SE | 0.5 | 0.75 | 0.25 | 0.25 | 0.25 | 0.25 | 0.25 | 1 | 1 | 1 | 1 |
| AN | 0.25 | 0.5 | 0.25 | 0.25 | 0.25 | 0.25 | 0.25 | 0.5 | 0.5 | 1 | 1 |

### Table S6: Species delimitation results

A) Scores for models identified under different delimitation methods

| **Delimitation method** | **Score (COI)** | **Score (16S)** | **Interpretation** |
| --- | --- | --- | --- |
| **ASAP** | 6.5 (lowest) | 6.5 (lowest = 3.5) | The lower the ASAP score, the better the model. For COI, we used results from the best model whereas for 16S, we used results from the second-best model (which has also been shown to produce high accuracy) |
| **PTP** | 0.63 (out of a possible 1) | 0.98 (out of a possible 1) | This is the Average Support Value (ASV), which indicates the level of convergence between the support value of each node (belonging to speciation/coalescent process) and the maximum likelihood estimate. |

B) ASAP species supports: The support value for each cluster represents the probability of members of that group belonging to the same species (0-1). The higher the probability, the greater the chance that members of the group are from the same species. NA refers to probability values that were not computed by the web server. Also note that we were unable to run ASAP on the entire 16S Scutigeromorph dataset, and instead present results for a subset of the data that includes our data- Thereuoneminae.

| **Species (COI)** | **Taxa (106)** | **Support** | **Species (16S)** | **Taxa (96)** | **Support** |
| --- | --- | --- | --- | --- | --- |
| 1 | *Sphendononema guildingii* DNA101630 |  | 1 | *Ballonema* sp. DNA106728 | 0.34 |
| 2 | *Sphendononema rugosa* DNA104623 |  |  | *Ballonema* sp. DNA106902 |  |
| 3 | *Scutigerina malagassa* DNA101591 |  |  | *Ballonema* sp. DNA106903 |  |
| 4 | *Scutigerina weberi* DNA100455 |  | 2 | *Lassophora nossibei* DNA102102 |  |
| 5 | *Scutigerina cf. weberi* DNA101590 |  | 3 | *Lassophora* sp. DNA106727 |  |
| 6 | *Scutigerina weberi* DNA106731 |  | 4 | *Allothereua bidenticulata* DNA101589 | NA |
| 7 | *Scutigerina weberi* DNA106733 |  |  | *Allothereua serrulata* DNA100262 |  |
| 8 | *Dendrothereua homa* DNA102576 |  | 5 | *Allothereua serrulata* DNA101045 |  |
| 9 | *Dendrothereua nubila* DNA101791 |  | 6 | *Allothereua linderi* DNA101463 |  |
| 10 | *Dendrothereua* sp. DNA106729 | NA | 7 | *Allothereua linderi* DNA101979 |  |
|  | *Dendrothereua* sp. DNA106730 |  | 8 | *Allothereua maculata* DNA101982 | 0.69 |
| 11 | *Dendrothereua* sp. DNA106854 |  |  | *Allothereua maculata* DNA101983 |  |
| 12 | *Dendrothereua* sp. IZ89420 |  |  | *Allothereua maculata* DNA101986 |  |
| 13 | *Dendrothereua* sp. IZ125268 |  |  | *Allothereua maculata* DNA101987 |  |
| 14 | *Scutigera coleoptrata* DNA100259 | NA |  | *Allothereua maculata* DNA101988 |  |
|  | *Scutigera coleoptrata* DNA102368 |  | 9 | *Parascutigera festiva* DNA102584 |  |
| 15 | *Ballonema* sp. DNA106728 | 0.96 | 10 | *Parascutigera guttata* DNA101971 | NA |
|  | *Ballonema* sp. DNA106902 |  |  | *Parascutigera guttata* DNA101973 |  |
|  | *Ballonema* sp. DNA106903 |  | 11 | *Parascutigera guttata* DNA102317 |  |
| 16 | *Lassophora nossibei* DNA102102 |  | 12 | *Parascutigera latericia* DNA101046 | NA |
| 17 | *Lassophora* sp. DNA106727 |  |  | *Parascutigera latericia* DNA102123 |  |
| 18 | *Allothereua bidenticulata* DNA101589 |  | 13 | *Parascutigera nubila* DNA103553 |  |
| 19 | *Allothereua linderi* DNA101463 |  | 14 | *Parascutigera* sp. QLD 1 DNA101974 |  |
| 20 | *Allothereua maculata* DNA101982 | 0.39 | 15 | *Parascutigera* sp. QLD 2 DNA101972 |  |
|  | *Allothereua maculata* DNA101983 |  | 16 | *Parascutigera* sp. QLD 3 DNA101977 | NA |
|  | *Allothereua maculata* DNA101987 |  |  | *Parascutigera* sp. QLD 3 DNA101978 |  |
|  | *Allothereua maculata* DNA101986 |  | 17 | *Parascutigera* cf. sphinx DNA101980 | NA |
|  | *Allothereua maculata* DNA101988 |  |  | *Parascutigera* cf. sphinx DNA101981 |  |
| 21 | *Allothereua serrulata* DNA100262 |  | 18 | *PIPlbarascutigera incola* DNA101997 |  |
| 22 | *Allothereua serrulata* DNA101045 |  | 19 | *Thereuopoda clunifera* DNA100260 |  |
| 23 | *Parascutigera festiva* DNA100635 |  | 20 | *Thereuopoda longicornis* DNA101461 |  |
| 24 | *Parascutigera festiva* DNA102584 |  | 21 | *Thereuonema tuberculata* DNA101632 | 0.88 |
| 25 | *Parascutigera guttata* DNA102317 |  |  | *Thereuonema turkestana* DNA101090 |  |
| 26 | *Parascutigera latericia* DNA101046 |  |  | *Thereuonema turkestana* DNA101091 |  |
| 27 | *Parascutigera latericia* DNA102123 |  | 22 | Thereuoneminae sp. IZ103188 |  |
| 28 | *Parascutigera latericia* DNA102124 | NA | 23 | Thereuoneminae sp. IZ103200 |  |
|  | *Parascutigera* sp. QLD 1 DNA101974 |  | 24 | *Thereuopodina* sp. DNA101462 |  |
| 29 | *Parascutigera nubila* DNA103553 |  | 25 | CCMB1125 | 0.98 |
| 30 | *Parascutigera* sp. QLD 3 DNA101977 |  |  | CCMB1126 |  |
| 31 | *Parascutigera* sp. QLD 3 DNA101978 |  |  | CCMB1069 |  |
| 32 | *Parascutigera* cf. sphinx DNA101980 | 0.21 |  | CCMB1098 |  |
|  | *Parascutigera* cf. sphinx DNA101981 |  |  | CCMB1003 |  |
|  | *Parascutigera* cf. sphinx DNA101985 |  |  | CCMB973 |  |
| 33 | *Thereuopoda clunifera* DNA100260 |  |  | CCMB1347 |  |
| 34 | *Thereuopoda longicornis* DNA101461 |  |  | CCMB1353 |  |
| 35 | *Thereuonema tuberculata* DNA101632 |  |  | CCMB2082 |  |
| 36 | *Thereuonema turkestana* DNA101090 | NA |  | CCMB1474 |  |
|  | *Thereuonema turkestana* DNA101091 |  |  | CCMB551 |  |
| 37 | Thereuoneminae sp. IZ103188 |  |  | CCMB615 |  |
| 38 | Thereuoneminae sp. IZ103200 |  |  | CCMB618 |  |
| 39 | CCMB2298 | NA |  | CCMB664 |  |
|  | CCMB2299 |  |  | CCMB552 |  |
| 40 | CCMB1125 | 0.75 |  | CCMB703 |  |
|  | CCMB1126 |  |  | CCMB702 |  |
|  | CCMB1069 |  |  | CCMB704 |  |
|  | CCMB1098 |  |  | CCMB705 |  |
|  | CCMB1003 |  |  | CCMB1534 |  |
| 41 | CCMB1347 | 0.91 |  | CCMB1644 |  |
|  | CCMB2030 |  |  | CCMB1663 |  |
|  | CCMB2032 |  |  | CCMB733 |  |
|  | CCMB2082 |  |  | CCMB1664 |  |
|  | CCMB1353 |  |  | CCMB717 |  |
|  | CCMB1474 |  |  | CCMB1673 |  |
|  | CCMB551 |  |  | CCMB392 |  |
|  | CCMB703 |  |  | CCMB466 |  |
|  | CCMB552 |  |  | CCMB443 |  |
|  | CCMB617 |  |  | CCMB485 |  |
|  | CCMB704 |  |  | CCMB747 |  |
|  | CCMB618 |  |  | CCMB897 |  |
|  | CCMB701 |  |  | CCMB899 |  |
|  | CCMB615 |  |  | CCMB880 |  |
|  | CCMB705 |  |  | CCMB939 |  |
|  | CCMB702 |  |  | CCMB749 |  |
|  | CCMB616 |  |  | CCMB1404 |  |
|  | CCMB664 |  |  | CCMB898 |  |
|  | CCMB661 |  |  | CES07221 |  |
|  | CCMB1534 |  | 26 | CCMB2029 | 0.25 |
|  | CCMB1591 |  |  | CCMB2031 |  |
|  | CCMB1644 |  |  | CCMB2073 |  |
|  | CCMB1663 |  |  | CCMB2126 |  |
|  | CCMB1664 |  | 27 | CCMB2227 | NA |
|  | CCMB1673 |  |  | CCMB2229 |  |
|  | CCMB733 |  | 28 | CCMB442 | NA |
|  | CCMB973 |  |  | CES1529 |  |
|  | CCMB392 |  | 29 | CCMB4070 | 0.99 |
|  | CCMB485 |  |  | CCMB100 |  |
|  | CCMB717 |  |  | CCMB167 |  |
|  | CCMB466 |  |  | CCMB4072 |  |
|  | CCMB443 |  |  | CCMB1202 |  |
| 42 | CCMB747 | 0.9 |  | CCMB1231 |  |
|  | CCMB897 |  |  | CCMB798 |  |
|  | CCMB899 |  |  | CCMB2230 |  |
|  | CCMB900 |  |  | CCMB2228 |  |
|  | CCMB880 |  | 30 | CCMB4071 |  |
|  | CCMB939 |  | 31 | CCMB10 |  |
| 43 | CCMB749 | NA | 32 | CES1518 |  |
|  | CCMB898 |  |  |  |  |
| 44 | CCMB2029 | NA |  |  |  |
|  | CCMB2031 |  |  |  |  |
| 45 | CCMB765 |  |  |  |  |
| 46 | CCMB442 |  |  |  |  |
| 47 | CCMB4072 |  |  |  |  |
| 48 | CCMB1404 |  |  |  |  |
| 49 | CCMB1918 |  |  |  |  |
| 50 | CCMB2227 |  |  |  |  |
| 51 | CCMB2229 |  |  |  |  |

C) PTP species supports: The support value for each cluster represents the proportion of times that node was considered part of the speciation process in the MCMC run. Hence, a value close to 1 would indicate that members of that group are likely to belong to different species (speciation process), whereas a value close to 0 would mean that members of the group are likely to belong to the same species (coalescent process).

| **Species (CO1)** | **Taxa (104)** | **Support** | **Species (16S)** | **Taxa (117)** | **Support** |
| --- | --- | --- | --- | --- | --- |
| 1 | *Scutigerina malagassa* DNA101591 |  | 1 | *Scutigerina hova* DNA101592 |  |
| 2 | *Scutigerina weberi* DNA100455 |  | 2 | *Scutigerina malagassa* DNA101591 |  |
| 3 | *Scutigerina cf. weberi* DNA101590 |  | 3 | *Scutigerina weberi* DNA100455 |  |
| 4 | *Scutigerina weberi* DNA106731 |  | 4 | *Scutigerina cf. weberi* DNA101590 |  |
| 5 | *Scutigerina weberi* DNA106733 |  | 5 | *Scutigerina weberi* DNA106731 |  |
| 6 | *Dendrothereua homa* DNA102576 |  | 6 | *Scutigerina weberi* DNA106732 |  |
| 7 | *Dendrothereua nubila* DNA101791 |  | 7 | *Dendrothereua homa* DNA102576 |  |
| 8 | *Dendrothereua* sp. DNA106729 |  | 8 | *Dendrothereua nubila* DNA101791 |  |
| 9 | *Dendrothereua* sp. DNA106730 |  | 9 | *Dendrothereua* sp. DNA106729 | 0 |
| 10 | *Dendrothereua* sp. DNA106854 |  |  | *Dendrothereua* sp. DNA106730 |  |
| 11 | *Dendrothereua* sp. IZ89420 |  | 10 | *Dendrothereua* sp. DNA107050 |  |
| 12 | *Dendrothereua* sp. IZ125268 |  | 11 | *Dendrothereua* sp. IZ89420 |  |
| 13 | *Scutigera coleoptrata* DNA100259 | 0 | 12 | *Scutigera coleoptrata* DNA100198 | 0 |
|  | *Scutigera coleoptrata* DNA102368 |  |  | *Scutigera coleoptrata* DNA100259 |  |
| 14 | *Ballonema* sp. DNA106728 |  |  | *Scutigera coleoptrata* DNA102327 |  |
| 15 | *Ballonema* sp. DNA106902 |  |  | *Scutigera coleoptrata* DNA102329 |  |
| 16 | *Ballonema* sp. DNA106903 |  |  | *Scutigera coleoptrata* DNA102367 |  |
| 17 | *Lassophora nossibei* DNA102102 |  |  | *Scutigera coleoptrata* DNA102368 |  |
| 18 | *Lassophora* sp. DNA106727 |  |  | *Scutigera coleoptrata* DNA102577 |  |
| 19 | *Allothereua bidenticulata* DNA101589 |  |  | *Scutigera coleoptrata* DNA1001980 |  |
| 20 | *Allothereua linderi* DNA101463 |  | 13 | *Ballonema* sp. DNA106728 | 0 |
| 21 | *Allothereua maculata* DNA101982 | 0 |  | *Ballonema* sp. DNA106902 |  |
|  | *Allothereua maculata* DNA101983 |  |  | *Ballonema* sp. DNA106903 |  |
|  | *Allothereua maculata* DNA101987 |  | 14 | *Lassophora nossibei* DNA102102 |  |
| 22 | *Allothereua maculata* DNA101986 | 0 | 15 | *Lassophora* sp. DNA106727 |  |
|  | *Allothereua maculata* DNA101988 |  | 16 | *Tachythereua* sp. DNA102575 |  |
| 23 | *Allothereua serrulata* DNA100262 |  | 17 | *Allothereua bidenticulata* DNA101589 |  |
| 24 | *Allothereua serrulata* DNA101045 |  | 18 | *Allothereua linderi* DNA101463 |  |
| 25 | *Parascutigera festiva* DNA100635 |  | 19 | *Allothereua linderi* DNA101979 |  |
| 26 | *Parascutigera festiva* DNA102584 |  | 20 | *Allothereua maculata* DNA101982 | 0 |
| 27 | *Parascutigera guttata* DNA102317 |  |  | *Allothereua maculata* DNA101983 |  |
| 28 | *Parascutigera latericia* DNA101046 |  |  | *Allothereua maculata* DNA101986 |  |
| 29 | *Parascutigera latericia* DNA102123 |  |  | *Allothereua maculata* DNA101987 |  |
| 30 | *Parascutigera latericia* DNA102124 | 0 |  | *Allothereua maculata* DNA101988 |  |
|  | *Parascutigera* sp. QLD 1 DNA101974 |  | 21 | *Allothereua serrulata* DNA100262 |  |
| 31 | *Parascutigera nubila* DNA103553 |  | 22 | *Allothereua serrulata* DNA101045 |  |
| 32 | *Parascutigera* sp. QLD 3 DNA101977 |  | 23 | *Parascutigera festiva* DNA102584 |  |
| 33 | *Parascutigera* sp. QLD 3 DNA101978 |  | 24 | *Parascutigera guttata* DNA101971 |  |
| 34 | *Parascutigera* cf. sphinx DNA101980 | 0 | 25 | *Parascutigera guttata* DNA101973 |  |
|  | *Parascutigera* cf. sphinx DNA101981 |  | 26 | *Parascutigera guttata* DNA102317 |  |
|  | *Parascutigera* cf. sphinx DNA101985 |  | 27 | *Parascutigera latericia* DNA101046 |  |
| 35 | *Thereuopoda clunifera* DNA100260 |  | 28 | *Parascutigera latericia* DNA102123 |  |
| 36 | *Thereuopoda longicornis* DNA101461 |  | 29 | *Parascutigera nubila* DNA103553 |  |
| 37 | *Thereuonema tuberculata* DNA101632 |  | 30 | *Parascutigera* sp. QLD 1 DNA101974 |  |
| 38 | *Thereuonema turkestana* DNA101090 |  | 31 | *Parascutigera* sp. QLD 2 DNA101972 |  |
| 39 | *Thereuonema turkestana* DNA101091 |  | 32 | *Parascutigera* sp. QLD 3 DNA101977 | 0 |
| 40 | Thereuoneminae sp. IZ103188 |  |  | *Parascutigera* sp. QLD 3 DNA101978 |  |
| 41 | Thereuoneminae sp. IZ103200 |  | 33 | *Parascutigera* cf. sphinx DNA101980 | 0 |
| 42 | CCMB2298 | 0 |  | *Parascutigera* cf. sphinx DNA101981 |  |
|  | CCMB2299 |  | 34 | *PIPlbarascutigera incola* DNA101997 |  |
| 43 | CCMB1125 | 0 | 35 | *Thereuopoda clunifera* DNA100260 |  |
|  | CCMB1126 |  | 36 | *Thereuopoda longicornis* DNA101461 |  |
|  | CCMB1003 |  | 37 | *Thereuonema tuberculata* DNA101632 |  |
| 44 | CCMB1069 | 0 | 38 | *Thereuonema turkestana* DNA101090 | 0 |
|  | CCMB1098 |  |  | *Thereuonema turkestana* DNA101091 |  |
| 45 | CCMB1347 | 0 | 39 | Thereuoneminae sp. IZ103188 |  |
|  | CCMB2030 |  | 40 | Thereuoneminae sp. IZ103200 |  |
|  | CCMB2032 |  | 41 | *Thereuopodina* sp. DNA101462 |  |
|  | CCMB2082 |  | 42 | CCMB1125 | 0 |
|  | CCMB1353 |  |  | CCMB1126 |  |
|  | CCMB1474 |  |  | CCMB1069 |  |
|  | CCMB551 |  |  | CCMB1098 |  |
|  | CCMB703 |  |  | CCMB1003 |  |
|  | CCMB552 |  |  | CCMB973 |  |
|  | CCMB617 |  |  | CCMB1347 |  |
|  | CCMB704 |  |  | CCMB1353 |  |
|  | CCMB618 |  |  | CCMB2082 |  |
|  | CCMB701 |  |  | CCMB1474 |  |
|  | CCMB615 |  |  | CCMB551 |  |
|  | CCMB705 |  |  | CCMB615 |  |
|  | CCMB702 |  |  | CCMB618 |  |
|  | CCMB616 |  |  | CCMB664 |  |
|  | CCMB664 |  |  | CCMB552 |  |
|  | CCMB661 |  |  | CCMB703 |  |
|  | CCMB1534 |  |  | CCMB702 |  |
|  | CCMB1591 |  |  | CCMB704 |  |
|  | CCMB1644 |  |  | CCMB705 |  |
|  | CCMB1663 |  |  | CCMB1534 |  |
|  | CCMB1664 |  |  | CCMB1644 |  |
|  | CCMB1673 |  |  | CCMB1663 |  |
|  | CCMB733 |  |  | CCMB733 |  |
|  | CCMB973 |  |  | CCMB1664 |  |
|  | CCMB392 |  |  | CCMB717 |  |
|  | CCMB466 |  |  | CCMB1673 |  |
|  | CCMB485 |  |  | CCMB392 |  |
|  | CCMB717 |  |  | CCMB466 |  |
|  | CCMB443 |  |  | CCMB443 |  |
| 46 | CCMB747 | 0 |  | CCMB485 |  |
|  | CCMB897 |  | 43 | CCMB747 | 0 |
|  | CCMB899 |  |  | CCMB897 |  |
|  | CCMB900 |  |  | CCMB899 |  |
|  | CCMB880 |  |  | CCMB880 |  |
|  | CCMB939 |  |  | CCMB939 |  |
| 47 | CCMB749 | 0 |  | CCMB749 |  |
|  | CCMB898 |  |  | CCMB898 |  |
| 48 | CCMB2029 | 0 |  | CES07221 |  |
|  | CCMB2031 |  | 44 | CCMB2029 | 0 |
| 49 | CCMB765 |  |  | CCMB2031 |  |
| 50 | CCMB442 |  |  | CCMB2073 |  |
| 51 | CCMB4072 |  |  | CCMB2126 |  |
| 52 | CCMB1404 |  | 45 | CCMB2227 | 0 |
| 53 | CCMB1918 |  |  | CCMB2229 |  |
| 54 | CCMB2227 |  | 46 | CCMB442 | 0 |
| 55 | CCMB2229 |  |  | CES1529 |  |
|  |  |  | 47 | CCMB4070 | 0 |
|  |  |  |  | CCMB100 |  |
|  |  |  |  | CCMB167 |  |
|  |  |  |  | CCMB4072 |  |
|  |  |  | 48 | CCMB1202 | 0 |
|  |  |  |  | CCMB1231 |  |
|  |  |  |  | CCMB798 |  |
|  |  |  | 49 | CCMB4071 |  |
|  |  |  | 50 | CCMB1404 |  |
|  |  |  | 51 | CCMB10 |  |
|  |  |  | 52 | CES1518 |  |
|  |  |  | 53 | CCMB2230 |  |
|  |  |  | 54 | CCMB2228 |  |

### Table S6: BSM event summaries

This table presents counts of evolutionary events across the Scutigeromorpha phylogeny estimated by Biogeographic Stochastic Mapping for 100 iterations on a DEC+J model. Note that range-switching is set to 0 under DEC and DEC+J models in BioGeoBEARS.

|  | **Founder speciation** | **Range switching** | **Anagenetic dispersal** | **Extinction** | **Subset sympatry** | **Vicariance** | **Sympatry** |
| --- | --- | --- | --- | --- | --- | --- | --- |
| Mean | 7.4 | 0 | 13.18 | 0 | 8.48 | 3.89 | 33.23 |
| SD | 2.1 | 0 | 2.03 | 0 | 2.01 | 1.7 | 1.48 |
